# Temporal and state abstractions for efficient learning, transfer and composition in humans

**DOI:** 10.1101/2020.02.20.958587

**Authors:** Liyu Xia, Anne G. E. Collins

**Affiliations:** University of California, Berkeley

**Keywords:** Hierarchical Reinforcement Learning, The Options Framework, Transfer Learning

## Abstract

Humans use prior knowledge to efficiently solve novel tasks, but how they structure past knowledge to enable such fast generalization is not well understood. We recently proposed that hierarchical state abstraction enabled generalization of simple one-step rules, by inferring context clusters for each rule. However, humans’ daily tasks are often temporally extended, and necessitate more complex multi-step, hierarchically structured strategies. The options framework in hierarchical reinforcement learning provides a theoretical framework for representing such transferable strategies. Options are abstract multi-step policies, assembled from simpler one-step actions or other options, that can represent meaningful reusable strategies as temporal abstractions. We developed a novel sequential decision making protocol to test if humans learn and transfer multi-step options. In a series of four experiments, we found transfer effects at multiple hierarchical levels of abstraction that could not be explained by flat reinforcement learning models or hierarchical models lacking temporal abstraction. We extended the options framework to develop a quantitative model that blends temporal and state abstractions. Our model captures the transfer effects observed in human participants. Our results provide evidence that humans create and compose hierarchical options, and use them to explore in novel contexts, consequently transferring past knowledge and speeding up learning.

## 1. Introduction

Recent advances have shown that reinforcement learning algorithms (RL, [1]) can give rise to extremely powerful artificial intelligence (AI) systems ([2, 3]). RL modeling has also greatly helped advance our understanding of human behavior ([4, 5, 6, 7, 8, 9]). However, despite tremendous recent progress, artificial RL agents are unable to mimic and capture humans’ ability to learn fast, efficiently, as well as transfer and generalize knowledge ([10, 11, 12]).

Human behavior and cognition possesses two key features that are essential to humans’ efficient and flexible learning: cognitive representations are hierarchical ([13, 14, 15, 16]) and compositional ([10]). Hierarchy has been identified as a crucial element of cognition in multiple domains such as perception ([17, 18, 19, 20]), decision making ([21, 22, 23, 16, 24, 25, 26, 27, 28, 29]), and learning [30, 31, 9, 32, 29]. Hierarchy in choices is often temporal ([33, 34]): choices may be described at multiple degrees of granularity by breaking them down into more and more basic chunks. For example, the task of making dinner can be broken down to making potatoes and making black beans; making potatoes can be broken down into sub-tasks such as cutting potatoes, roasting, etc. However, hierarchical levels may also represent different degrees of state abstractions at a similar time scale([14, 16, 9, 35]): for example, you may decide to make dinner (highest, most abstract level), which will consist of a salad, which will specifically be a Cesar salad (lowest, most concrete level).

Human behavior is also compositional: humans are able to compose simpler skills together in novel ways to solve new tasks in real life. For example, we can combine cutting potatoes with different routines to accomplish various tasks including fried potatoes, meshed potatoes, etc. Compositionality goes hand in hand with hierarchy, as it assumes the existence of different levels of skills. It has also been central to the study of human cognition ([36, 37, 38]) and artificial agents ([39, 40, 41, 42]).

The hierarchical reinforcement learning (HRL) options framework [43], originally proposed in AI, incorporates both hierarchy and compositionality features in an effort to make learning more flexible and efficient. The options framework augments traditional RL algorithms with temporal abstractions called options. Broadly summarized, options are temporally-extended multistep policies assembled from simple actions or other options to achieve a meaningful subgoal (see [43] for a formal definition). Consider making potatoes as an example option. We can break down the task into sub-options such as cutting potatoes, roasting, etc. These sub-options can be further divided into simpler tasks. In the HRL options framework, agents can learn option-specific policies (e.g. how to make potatoes) by using, for example, subgoals as pseudo-rewards that reinforce within-option choices. Options are referred to as *temporal abstractions* because selecting an option is a single decision step, but this single decision may itself contain a series of decisions, so that time is compressed in a single decision.

Each option is additionally characterized by an initiation set (the set of states where the option can be initiated), and a termination function that maps each state to the probability of terminating the current option. For example, the initiation set for the option of making potatoes might be kitchen, and the option might terminate when the potatoes are cooked. Agents can also learn when to select options (e.g. make potatoes for breakfast in the US, but not in France) by using normal reinforcement signals.

The options framework provides many theoretical benefits for learning ([11, 44]), assuming that useful options are available. Unlike traditional RL algorithms that only learn step-by-step policies, options help explore more efficiently and plan longer term. For example, when we learn how to cook a new kind of potato, we already know how to cut potatoes. Moreover, we can plan with high-level behavioral modules such as cutting potatoes, instead of planning in terms of reaching, grabbing, and peeling. If non-useful options are available, the options framework predicts that learning is instead slowed down [11]. The question of how to identify and create useful options has been a topic of active and intense research in AI ([45, 46, 47, 48, 49, 50, 51, 52, 53, 54]).

Note that the options framework is not the first attempt to incorporate hierarchy and compositionality to model complex human cognition. Within psychology in particular, “option” echoes the idea of “chunking” in cognitive architecture literature ([55, 56]). However, one distinct aspect of the options framework is its objective of reward maximization ([11]), which is naturally inherited as an augmentation of traditional flat RL (although see ([57, 58]) for initial work on combining ideas from reward maximization of RL with cognitive architectures). Importantly, this objective of reward maximization has proven to be relevant and instrumental in revealing neural mechanisms underlying learning and adaptation ([59]).

Moreover, recent literature ([12, 60, 61, 62, 63]) provides behavioral and neural support for options as a useful model of human learning and decision making. [12, 63] showed that participants were able to spontaneously identify bottleneck states from transition statistics, which aligned with graph-theoretic objectives for option discovery developed in AI ([46]). In addition, in hierarchical decision-making tasks, [60, 61, 62] showed that human participants signaled reward prediction error (RPE), a key construct for RL algorithms, for both subgoals and overall goals. These results indicate that humans are able to identify meaningful subgoals, and to track sub-task progression, both key features of the options framework. [64, 65] have also suggested potential neural correlates to implementing the computations required to use options.

However, the fundamental question of whether and how humans learn and use options during learning remains unanswered ([12]): there is little work probing the learning dynamics in tasks with a temporal hierarchy, or directly testing the theoretical benefits of options in a behavioral setting. In particular, do humans create options in such a way that they can flexibly reuse them in new problems? If so, how flexible is this transfer? Previous research ([9, 32, 66]) showed evidence for flexible creation and transfer of a simple type of options that operate in non-sequential environments: one-step policies, also called task-sets ([67]). [9, 32, 66] showed that humans can create multiple task-sets over the same state space in a context-dependent manner in a contextual multi-armed bandit task. Furthermore, humans can cluster different contexts together if the task-set is successful. This clustering structure provides opportunities for transfer, since anything newly learned for one of the contexts can be immediately generalized to all the others in the same cluster. Moreover, human participants can identify novel contexts as part of an existing cluster if the cluster-defined strategy proves successful, resulting in more efficient exploration and faster learning.

However, the task-sets framework only supports hierarchy in “state/action space abstraction”, not hierarchical structure in time (also called “temporal abstraction”), an essential component of the options framework. Here, we propose that combining state abstraction from task-set transfer ([9, 32, 66]) and temporal abstraction from the options framework ([43]) can provide important insights into complex human cognition. The additional temporal hierarchical structure offered by options should enable transfer of prior knowledge at multiple levels of hierarchy, providing rich opportunity for capturing the flexibility of human transfer. For example, if humans have learned the simple sub-option of boiling water while learning how to make coffee, they do not need to re-learn it for learning to make tea or steamed potatoes; this sub-option can instead be naturally incorporated into a tea-making option, speeding up learning.

In this paper, we present a new experimental protocol that allows us to test whether humans create options when learning, and whether they use them in new contexts to explore more efficiently and transfer learned skills, at multiple levels of hierarchy. Our new two-stage learning game provides participants opportunities to create and transfer options at multiple levels of complexity. We also present a formal computational model that brings together aspects of the classic hierarchical RL options framework with the task-set model’s clustering and transfer Bayesian inference mechanisms. The model combines the benefits of both frameworks and makes specific predictions about option learning, transfer and exploration. Given that humans can transfer task-sets to novel contexts ([9, 32, 66]), we hypothesized that humans would learn and transfer options to guide exploration and achieve better learning performance, as captured by the model.Results of four experiments (3 replicated in an independent sample), testing different predictions in the same framework, showed that human participants are able to learn, flexibly transfer and compose options at multiple levels. Our computational model captured the observed patterns of behavior, supporting the importance of hierarchical representations of choices for flexible, efficient, generalizable learning and exploration.

## 2. Experiment 1

Experiment 1 was designed to test if human participants are able to learn and flexibly transfer options. We designed a sequential 2-step decision-making paradigm (where each step was a contextual 4-armed bandit) to allow participants to learn options at multiple levels of complexities. Options changed between blocks, but the design provided participants with opportunities to practice reusing previously learned options. In two final test blocks, we directly tested creation and transfer of options by changing and/or combining previously learned options in novel ways.

### 2.1. Methods

#### 2.1.1. Participants

All experiments were approved by the Institutional Review Board of the University of California, Berkeley. Experiment 1 was administered in-lab to UC Berkeley undergraduates who received course credit for their participation. 34 (22 female; age: mean = 20.6, sd = 1.6, min = 18, max = 24) UC Berkeley undergraduates participated in Experiment 1, and 9 participants were excluded due to incomplete data or poor learning performance (see results), resulting in 25 participants for data analysis.

For replication purposes, we also recruited participants through Amazon Mechanical Turk (MTurk) who performed the same experiment online. Participants were compensated a minimum of $3 per hour for their participation, with a bonus depending on their performance to incentivize them. 116 participants (65 female; see age range distribution in Table 3) finished the experiment. 61 participants were further excluded due to poor performance (see Sec 2.1.4), resulting in 55 participants for data analysis.

**Table 1:** Parameters for the main text.

| Exp | Sample | Model | $\alpha^1$ | $\beta^1$ | $\gamma^1$ | $f^1$ | $\alpha^2$ | $\beta^2$ | $\gamma^2$ | $f^2$ | m |
| --- | --- | --- | --- | --- | --- | --- | --- | --- | --- | --- | --- |
| Exp 1 | In-lab | Naive | 0.5 | 4 | NA | 0.0025 | 0.7 | 10 | NA | 0.0001 | 0.01 |
|  |  | Flat | 0.5 | 4 | NA | 0.0025 | 0.7 | 10 | NA | 0.0001 | 0.01 |
|  |  | Task-Set | 1 | 2 | 14 | 0.0004 | 0.8 | 3 | 3 | 0.0002 | 0.01 |
|  |  | Option | 1 | 2 | 14 | 0.0004 | 0.8 | 3 | 3 | 0.0002 | 0.01 |
|  | Mturk | Option | 0.8 | 3 | 100 | 0.01 | 0.6 | 6 | 5 | 0.004 | 0.01 |
| Exp 2 | In-lab | Option | 0.7 | 3 | 13 | 0.001 | 0.6 | 4 | 5 | 0.001 | 0.01 |
| Exp 3 | In-lab | Option | 0.7 | 4 | 100 | 0.01 | 0.8 | 5 | 15 | 0.001 | 0.01 |
|  | Mturk | Option | 0.7 | 4 | 100 | 0.01 | 0.8 | 5 | 15 | 0.005 | 0.01 |
| Exp 4 | In-lab | Option | 0.6 | 4 | 100 | 0.01 | 0.8 | 5 | 4 | 0.0002 | 0.01 |
|  | Mturk | Option | 0.6 | 4 | 100 | 0.01 | 0.4 | 4 | 5 | 0.002 | 0.01 |
|  |  | Task-Set | 0.6 | 4 | 100 | 0.01 | 0.4 | 4 | 5 | 0.002 | 0.01 |

**Table 2:** A second set of parameters that is constrained but still replicate transfer effects qualitatively.

| Exp | Sample | Model | $\alpha^1$ | $\beta^1$ | $\gamma^1$ | $f^1$ | $\alpha^2$ | $\beta^2$ | $\gamma^2$ | $f^2$ | m |
| --- | --- | --- | --- | --- | --- | --- | --- | --- | --- | --- | --- |
| Exp 1 | In-lab | Naive | 0.7 | 4 | NA | 0.001 | 0.7 | 4 | NA | 0.001 | 0.01 |
|  |  | Flat | 0.7 | 4 | NA | 0.001 | 0.7 | 4 | NA | 0.001 | 0.01 |
|  |  | Task-Set | 0.7 | 4 | 14 | 0.001 | 0.7 | 4 | 4 | 0.001 | 0.01 |
|  |  | Option | 0.7 | 4 | 14 | 0.001 | 0.7 | 4 | 4 | 0.001 | 0.01 |
|  | Mturk | Option | 0.7 | 4 | 100 | 0.01 | 0.5 | 4 | 4 | 0.005 | 0.01 |
| Exp 2 | In-lab | Option | 0.7 | 4 | 100 | 0.01 | 0.7 | 4 | 4 | 0.001 | 0.01 |
| Exp 3 | In-lab | Option | 0.7 | 4 | 100 | 0.01 | 0.7 | 4 | 20 | 0.001 | 0.01 |
|  | Mturk | Option | 0.7 | 4 | 100 | 0.01 | 0.5 | 4 | 20 | 0.005 | 0.01 |
| Exp 4 | In-lab | Option | 0.7 | 4 | 100 | 0.01 | 0.7 | 4 | 4 | 0.001 | 0.01 |
|  | Mturk | Option | 0.7 | 4 | 100 | 0.01 | 0.5 | 4 | 4 | 0.005 | 0.01 |

**Table 3:** Age range distribution for Mturk participants in Experiments 1, 3, and 4.

| Exp | 18-25 | 26-30 | 31-35 | 36-40 | 41+ | Unknown | Total |
| --- | --- | --- | --- | --- | --- | --- | --- |
| Exp 1 | 14 | 18 | 26 | 23 | 33 | 2 | 116 |
| Exp 3 | 4 | 9 | 18 | 9 | 25 | 0 | 65 |
| Exp 4 | 14 | 17 | 24 | 15 | 40 | 0 | 110 |

#### 2.1.2. Experiment 1 in-lab Protocol

Experiment 1 consisted of eight 60-trial blocks (Fig. 1), with optional 20-second breaks in between blocks. In each block, the participants used deterministic truthful feedback to learn which of four keys to press for four different shapes. Each trial included two stages; each stage involved participants making choices in response to a single stimulus (Fig. 1A) by pressing one of four keys. Each trial started with one of two possible stimuli, henceforth the first stage stimuli (e.g. circle or square). Participants had 2 seconds to make a choice. Participants only moved on to the second stage of the trial when they pressed the correct key for the first stage stimulus, or after 10 unsuccessful key presses, which enabled them to potentially try all four keys for a given stimulus in a single trial. Successful key press for the first stage of a trial did not result in reward feedback, but triggered a transition to the second stage, where participants saw one of the two other stimuli, hence-forth labeled second stage stimuli (e.g. diamond and triangle). Both first stage stimuli led to both second stage stimuli equally often, and shapes were randomly assigned to either first or second stage across participants. In the second stage, participants also could not move on until they selected the correct choice (or selected wrong 10 times in a row for the same image). Participants received explicit feedback after each second stage choice: the screen indicated 1/0 point for pressing the correct/incorrect key, displayed for 0.5 second (Fig. 1A). After a correct second stage choice, participants saw a fixation cross for 0.5 second, followed by the next trial’s first stage stimulus. Each block contained 60 trials, with each first stage stimulus leading to each second stage stimulus 15 times in a pseudo-randomized sequence of trials.

**Figure 1:**
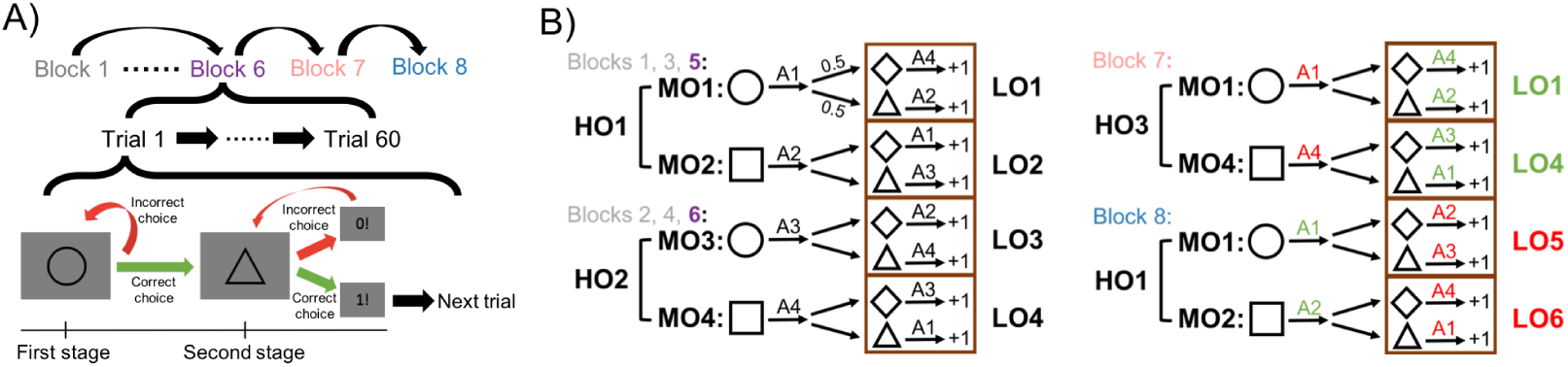
Experiment 1 protocol. (A) Block and trial structure: Blocks 1-6 were learning blocks, followed by two testing blocks: Blocks 7 and 8. Each block had 60 trials. In each trial, participants needed to select the correct response for the first stage stimulus (e.g. circle) in order to move on to the second stage stimulus (e.g. triangle), where they could win points by selecting the correct response. (B) Stimulus-action assignments: In Blocks 1-6, participants had the opportunity to learn options (extended policies) at three levels of complexity: high, middle, and low-level options (*HO, MO*, and *LO*). In the testing phase, Block 7 tested participants’ ability to reuse *MO* policies outside of their *HO* context, potentially eliciting positive transfer (green) of *LO*s in the second stage, and negative transfer (red) of choices in the first stage. Block 8 tested predicted positive transfer in the first stage, but negative transfer of *MO* policies in the second stage, by replacing old *LO*s by new ones. Blocks were color coded for later result figures: Blocks 1-4 gray; Blocks 5-6 purple; Block 7 rose; Block 8 blue.

Crucially, the correct stimulus-action assignments were designed to allow for the creation of multi-step policies and to test their grouping into sets of policies at multiple levels. In particular, second stage correct choices were dependent on what the first stage stimulus was. This encouraged participants to make temporally extended choices (potentially options): their second stage strategies needed to depend on the first stage. Assignments, illustrated in Fig. 1B, changed across blocks. Blocks 1, 3, 5 shared the same assignments; Blocks 2, 4, 6 shared the same assignments; this encouraged participants to not unlearn policies, but rather discover that they could reuse previously learned multi-level policies as a whole in new blocks.

Assignments in Blocks 7 and 8 intermixed some of the learning blocks assignments with new ones to test (positive and negative) transfer of options at various hierarchy levels. Specifically, the protocol was set up so that participants could learn up to 3 levels of hierarchical task structure (low, mid, and high level policies). More precisely, low-level options (*LO*) corresponded to second stage policies (a pair of stimulus-action associations, commonly labelled a *task-set*) ([67]). Mid-level options (*MO*) were policies over both first and second stage stimuli. High-level options (*HO*) were policies over *MO*’s (a pair of stimulus-*MO* associations in the first stage, which could be thought of as a *task-set over options*). As a concrete analogy, in Blocks 1, 3, 5, the participants learned how to make breakfast (*HO*_1_), consisting of potatoes (*MO*_1_) and eggs (*MO*_2_). Making potatoes (*MO*_1_) was broken down into cutting potatoes (the first stage) and then roasting (the second stage, *LO*_1_). In Blocks 2, 4, 6, participants learned how to make lunch (*HO*_2_), consisting of vegetables (*MO*_3_) and sandwich (*MO*_4_). Making vegetables (*MO*_3_) was broken down into combining vegetables (the first stage) and then steaming (the second stage, *LO*_3_).

Block 7 tested positive transfer of second stage policies and negative transfer of first stage policies. In particular, we combined the policies for potatoes from breakfast (*MO*_1_) and sandwich from lunch (*MO*_4_) to form a new policy *HO*_3_ (dinner). If participants build three levels of options, we expect positive transfer of mid-level options *MO*_1_ and *MO*_4_: participants should be unimpaired in making potatoes or a sandwich. However, we expect negative transfer of high-level options *HO*_1_ and *HO*_2_: participants seeing that making potatoes was rewarded might start making eggs as usual, instead of sandwich as rewarded here.

Block 8 tested positive transfer of first stage policies and negative transfer of second stage policies. In particular, the first stage of Block 8 shared the same assignments as Blocks 1, 3, 5 in the first stage, allowing participants to immediately transfer *HO*_1_. However, the second stage policies (*LO*_5_ and *LO*_6_) were novel, which might potentially result in negative transfer: for example, participants might try to transfer *LO*_1_ (roasting) following *MO*_1_ (make potatoes), but the second stage policy was changed to *LO*_5_ (e.g. frying).

#### 2.1.3. Experiment 1 MTurk Protocol

To replicate our findings, we ran a minimally modified version of Experiment 1 online via MTurk. The task was slightly shortened, due to evidence that in-lab participants reached asymptotic behavior (Supplementary Fig. S11) early in a block, and to make the experiment more acceptable to online workers. Blocks 1 and 2 had a minimum of 32 and a maximum of 60 trials, but participants moved on to the next block as soon as they reached a criterion of less than 1.5 key presses per second stage trial in the last 10 trials (the 55 Mturk participants included for data analysis on average used 42 (SD = 10, median = 37, min = 32, max = 60) trials in Block 1 and 39 (SD = 10, median = 33, min = 32, max = 60) trials in Block 2). Blocks 3-8 were all shortened to 32 trials, with each first stage stimulus leading to each second stage stimulus 8 times.

#### 2.1.4. Data analysis

We used the number of key presses until correct choice in each stage of a trial as an index of performance. Since the experiment would not progress unless the participants chose the correct action, more key presses indicates worse performance. Ceiling performance was 1 press per stage within a trial. Chance level was 2.5, assuming choosing 1 out of 4 keys randomly, unless indicated otherwise. To probe for any potential transfer effects, we calculated the average number of key presses at the beginning of each block (trials 1-10), before learning has saturated. As a stronger test of option transfer, we also calculated the probability that the first press for a given stimulus at each stage of a trial was correct in different blocks.

To rule out participants who were not engaged in the task, we excluded any participant who did not complete Blocks 5-8 within an allotted amount of time (6 minutes each) - indeed this could only happen if participants often reached the 10 key presses needed to move on to the next stage without the correct answer, a clear sign of no engagement.

We additionally excluded any participant whose average performance in the last 10 trials of either first or second stage in either Block 5 or 6 was at or below chance, since it indicated a lack of learning and engagement in both stages of the task. These exclusion criteria were applied to all experiments, including Mturk participants. Note that among 116 Mturk participants in Experiment 1, 104 were above chance in the second stage (the more difficult one), but only 55 were above chance in the first stage (the easier one). Thus most participants were excluded due to the first stage performance criterion. The same trend was true for the other two Mturk experiments: most Mturk participants were excluded due to performance in the first stage in Experiment 3 and Experiment 4. We hypothesize that the poor first stage performance in many is due to the task’s incentive structure - participants knew they only earned points (which were converted to monetary bonus for MTurk participants) in the second stage. All results were qualitatively similar to the ones reported in this paper for all experiments when we relaxed the exclusion criterion to include participants at chance in the first stage.

The options framework makes predictions about the specific choices made in response to a stimulus, beyond whether a choice is correct: the nature of the errors made can be informative ([9]). We categorized the specific choices participants made into meaningful choice types, to further test our predictions about potential option transfer effects. As the choice types were stage and experiment dependent, we describe the choice type definitions in the result sections where necessary. When performing choice type analysis, We only considered the first key press of the first or second stage in each trial to reduce noise. We also compared reaction time of difference choice types to test potential sequence learning effects.

For statistical testing, we used parametric tests (ANOVAs and paired t-test) when normality assumptions held, and non-parametric tests (Kruskall-Wallis and sign test) otherwise.

#### 2.1.5. Computational modeling

To quantitatively formalize our predictions, we designed a computational model for learning and transferring options, inspired by the classic HRL framework as well as other hierarchical RL literature [9, 43]. We simulated this model, as well as three other learning models that embody different hypotheses about learning in this task, to compare which model best captures patterns of human learning and transfer. All models were simulated 500 times. We did not fit the model to the trial-by-trial choices of participants: computing the likelihood of the hierarchical models is intractable, because we only observed the key presses, but not the choice of options. All results presented in the main text figures were simulated with parameters chosen to match participants’ behavioral patterns qualitatively and quantitatively well (Table 1). However, our qualitative predictions are largely independent of specific model parameters: we show in the supplement (Sec. 9.3) that a single set of parameters (Table 2), consistent across all experiments, makes the same qualitative predictions regarding transfer effects.

##### 2.1.5.1. The Naive Flat Model

The Naive Flat Model is a classic reinforcement learning model that learns Q-values to guide action selection in response to stimuli. In the first stage, it learns a Q-value table 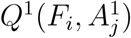, where *F*_1_ and *F*_2_ are two first stage stimuli, *A*_1_, …, *A*_4_ are four possible actions. We use superscript to index stage (1 means first stage, 2 means second stage). The Q-values are initialized to uninformative Q-values 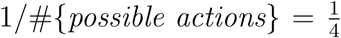. On each choice, a first stage policy is computed based on the first stage stimulus, *F*_*i*_, with the softmax function:

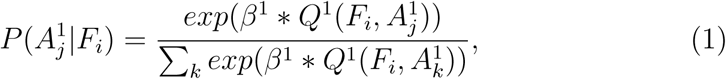

where *β*^1^ is the inverse temperature parameter. A first stage action *A*^1^, ranging from *A*_1_ to *A*_4_, is then sampled from this softmax policy. After observing the outcome (moving on to the second stage or not), the Q-values is updated with Q-learning ([1]):

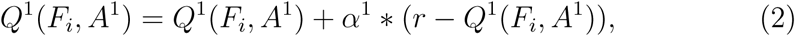

where *α*^1^ is the learning rate parameter, and *r* is 1 if *A*^1^ is correct and 0 otherwise.

In the second stage, the model similarly learns another Q-value table 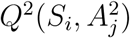, where *S*_1_ and *S*_2_ are two second stage stimuli, with learning rate *alpha*^2^ and inverse temperature *β*^2^. Note that it disregards the non-Markovian nature of the task: it learns the Q-values for the two second stage stimuli without remembering the first stage stimulus. As such, this model is a straw man model that cannot perform the task accurately, but exemplifies the limitations of classic RL in more realistic tasks, and serves as a benchmark.

At the start of a new block, the Naive Flat Model resets all Q-values to 1*/*4, and thus has to re-learn all Q-values from scratch. To better account for human behavior, we also included two forgetting parameters, *f* ^1^ and *f* ^2^. After each choice, the model decays all Q-values for the first stage based on *f* ^1^:

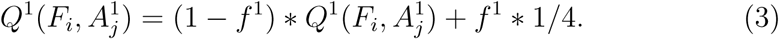

Forgetting in the second stage is implemented similarly.

Participants very quickly learned that the correct second stage action was different from the first stage one (see results). To account for this meta-learning heuristic, we add a meta-learning parameter *m* that discourages selecting the same action in the second stage as in the first stage. Specifically, if *π* is the second stage policy as computed from softmax, we set *P* (*A*^1^|*S*_*i*_) = *m*, where *A*^1^ is the action chosen in the first stage, and re-normalize:

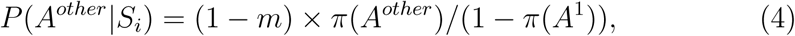

where *A*^*other*^ is any action other than *A*^1^.

Parameters *f* ^1^, *f* ^2^ and *m*, which capture memory mechanisms and heuristics orthogonal to option learning, are included in all models and implemented in the same way. In total, the Naive Flat Model has 7 parameters: *α*^1^, *β*^1^, *f* ^1^, *α*^2^, *β*^2^, *f* ^2^, *m*.

##### 2.1.5.2. The Flat Model

The Flat Model extends the Naive Flat Model with a single addition of first-stage memory, which makes this model able to perform the task well in both stages. Specifically, in the second stage, the Flat Model remembers the first stage stimulus by treating each of the 4 combinations of the first and second stage stimuli as a distinct state and learns Q-values for all 4 combinations. The Flat Model has the same 7 parameters as the Naive Flat Model.

##### 2.1.5.3. The Task-Set Model

The Task-Set Model is given the capability of transferring previously learned task-sets (one-step policies) with Bayesian inference. In the first stage, the model tracks the probability *P* ^1^ of selecting each first stage task-set *HO*_*i*_ in different first stage contexts 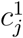, which encodes the current temporal (block) context (e.g. 8 contexts in the first stage of Experiment 1). In particular, the model uses a Chinese Restaurant Process (CRP) prior to select *HO* ([68]): if contexts 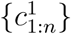 are clustered on *N* ^1^ ≤ *n HO*′*s*, when the model encounters a new context 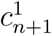, the prior probability of selecting a new high-level option *HO*_*n*+1_ in this new context is set to:

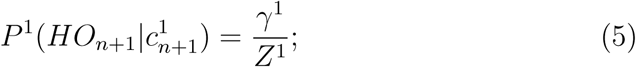

and the probability of reusing a previously created high-level option *HO*_*i*_ is set to:

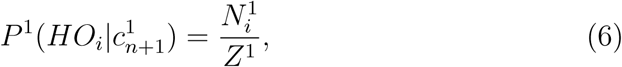

where *γ*^1^ is the clustering coefficient for the CRP, 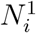 is the number of first stage contexts clustered on *HO*_*i*_, and 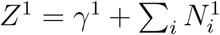 is the normalization constant. The new *HO*_*n*+1_ policy is initialized with uninformative Q-values 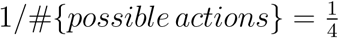. The model samples *HO* based on the conditional distribution over all *HO*’S given the current temporal context. The model also tracks *HO*-specific policies via Q-learning. Once an *HO* is selected, a first stage policy is computed based on the *HO*’s Q-values and the first stage stimulus *F*_*i*_ with softmax:

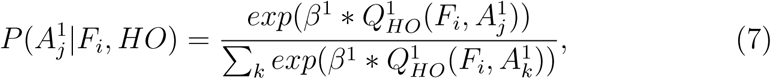

where *β*^1^ is the inverse temperature. A first stage action *A*^1^, ranging from *A*_1_ to *A*_4_, is then sampled from this softmax policy. After observing the outcome (moving on to the second stage or not), the model uses Bayes’ Theorem to update *P* ^1^:

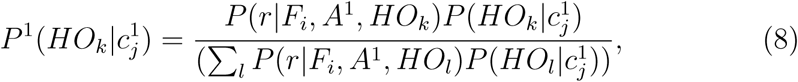

where *r* is 1 if *A*^1^ is correct and 0 otherwise, and 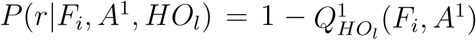 if *r* = 0, or 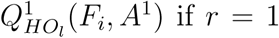 if *r* = 1. Then the Q-values of the *HO* with the highest posterior probability is updated:

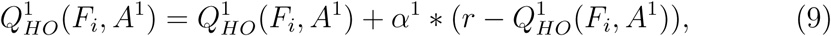

where *α*^1^ is the learning rate.

The second stage runs a separate CRP with *P* ^2^, similar to *P* ^1^ in the first stage, which guides selection of task-sets *LO* over second stage stimuli. All other are identical to the first stage except that the second stage contexts are determined by both temporal (block) context and the first stage stimulus (e.g. 16 contexts in the second stage of Experiment 1). All the equations of CRP, action selection and Q-learning remain the same. The Task-Set Model has 9 parameters: *α*^1^, *β*^1^, *γ*^1^, *f* ^1^, *α*^2^, *β*^2^, *γ*^2^, *f* ^2^, *m*.

##### 2.1.5.4. The Option Model

The Option Model extends the task-set model to include multi-step decisions (options *MO*). The first stage is identical to the Task-Set Model. However, in addition to just choosing an action, an *MO* is also activated. To simplify credit assignment, we assumed that selecting an action in the first stage is equivalent to selecting an *MO* as a whole: for example, selecting *A*_1_ for the circle activates *MO*_1_ (Fig. 1B).

The second stage is the same as the Task-Set Model, except that each *MO* has an *MO*-specific probability table 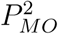. In the Task-Set Model, the CRP in the second stage using *P* ^2^ is independent of the first stage choices. In contrast, in the Option Model, the first stage choice determine which *MO* is activated, which then determines which probability table, 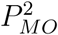, to use for running the CRP in the second stage. This implementation captures the essence of options in the HRL framework, in that selection of *MO* in the first stage constrains the policy chosen until the end of the second stage (where the option terminates). The Option Model has the same 9 parameters as the Task-Set Model.

### 2.2. Experiment 1 Results

#### 2.2.1. Participants do not use flat RL

Participants’ performance improved over Blocks 1-6 (Fig. 2A) and within blocks (Supplementary Fig. S11). This improvement may reflect the usual process of learning the task observed in most cognitive experiments, as indicated by the improvement between Block 1 and 2 (paired t-test, first stage: *t*(26) = 2.2, *p* = 0.03; second stage: *t*(26) = 3.9, *p* = 0.0006). However, it could also reflect participants’ ability to create options at three different levels in Blocks 1 and 2, and to successfully reuse them in Blocks 3-6 to adapt to changes in contingencies more efficiently. Below, we present specific analyses to probe option creation in test blocks. We used participants’ performance averaged over Blocks 5 and 6 as a benchmark for comparing against performance in test Blocks 7 and 8.

**Figure 2:**
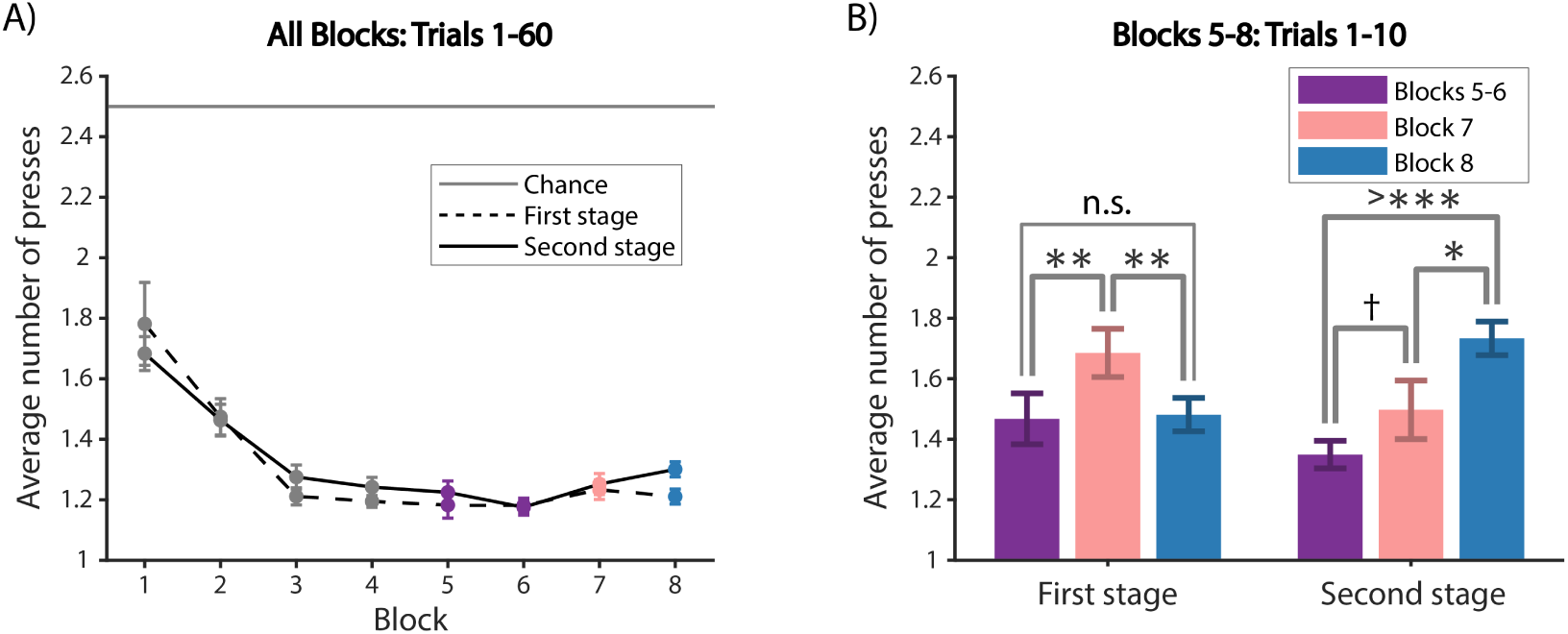
Experiment 1 general behavior. (A) Average number of key presses in the first and the second stages per block. Chance is 2.5, ceiling is 1 press. (B) Average number of key presses for the first 10 trials of Blocks 5-8 for the first (left) and second stages (right). We use n.s. to indicate *p* ≥ 0.1; † for *p* < 0.1; ∗ for *p* < 0.05; ∗∗ for *p* < 0.01; ∗∗∗ for *p* < 0.001; and >∗∗∗ for *p* < 0.0001. We indicated all statistical significance with these notations from now on.

We probed potential option transfer effects over the first 10 trials for each block (Fig. 2B), before behavior reached asymptote (Supplementary Fig. S11). In the first stage, there was a main effect of block on number of key presses (1-way repeated measure ANOVA, *F*(2, 48) = 6.9, *p* = 0.002). Specifically, participants pressed significantly more times in Block 7 than Blocks 5-6 and Block 8 (paired t-test, Blocks 5-6: *t*(24) = 3.0, *p* = 0.006; Block 8: *t*(24) = 3.0, *p* = 0.006). We also found no significant difference between the performance of circle and square in Block 7 (9.1). These results provide preliminary evidence for negative transfer of previously learned *HO* in Block 7: participants might attempt to reuse *HO*_1_ or *HO*_2_, since either policy is successful for half the trials, but is incorrect and thus results in more key presses in the first stage for the other half of the trials. There was no significant difference between Block 8 and Blocks 5-6 (paired t-test, *t*(24) = 0.25, *p* = 0.81). This provides initial evidence for positive transfer of *HO*_1_ in Block 8, since performance in the first stage of Block 8 was on par with Blocks 5-6.

In the second stage (Fig. 2B), there was also a main effect of block in number of key presses (1-way repeated measure ANOVA, *F*(2, 48) = 11, *p* < 0.0001). Specifically, participants pressed significantly more times in Block 8 than Block 7 and Blocks 5-6 (paired t-test, Block 7: *t*(24) = 2.4, *p* = 0.025; Blocks 5-6: *t*(24) = 5.8, *p* < 0.0001). The difference between Block 7 and Blocks 5-6 was marginally significant (paired t-test, *t*(24) = 2.0, *p* = 0.06). These results suggest participants positively transferred *MO* in the second stage of Block 7, where such generalization was helpful, since their performance was nearly not impaired compared to Blocks 5-6 where participants were able to reuse full *HO*. Furthermore, it suggests that they negatively transferred *MO* in the second stage of Block 8, where the first stage choice that respected the current *MO* was followed by a new *LO* for correct performance, and thus necessitated to create a new *MO*.

Behavioral results in both the first and second stages provide initial evidence for option learning and transfer at distinct levels, both positive – when previous policies can be helpfully reused – and negative – when they impair learning. To further validate our hypothesis that participants learned options, we compared the simulations of four models with human behavior (Table 1).

Among the four models (Fig. 3), only the Option Model and the Task-Set Model could account for the results. The Naive Flat Model could not achieve reasonable performance in the second stage because it ignored the non-Markovian aspect of the task - it was unable to learn two different sets of correct choices for a given second stage stimulus, because this required conditioning on the first stage stimulus (Fig. 1B). Thus, it serves to illustrate the limitations of classic RL, but is a straw man model in this task. The Flat Model achieved reasonable performance in both the first and second stages, being able to take into account the first stage in second stage decisions, but did not demonstrate any transfer effects. Thus, results so far replicate previous findings that participants create one-step policies or task-sets, that they can reuse in new contexts, leading to positive and negative transfer [9, 32, 66]. We now present new analyses to show that the findings extend to creating multi-step policies or options.

**Figure 3:**
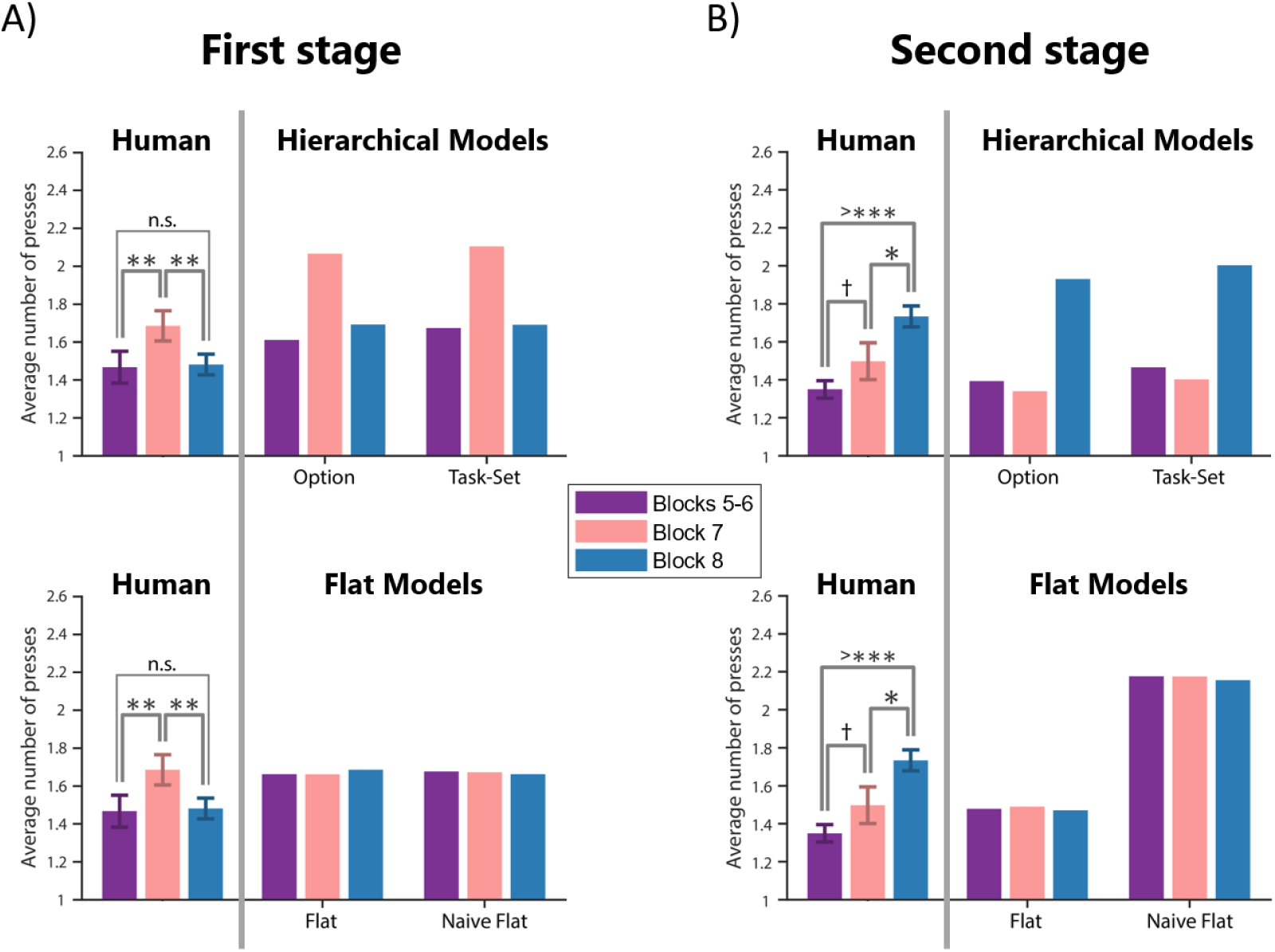
Experiment 1 transfer effects. Average number of first (A) and second (B) stage key presses in the first 10 trials of Block 5-8 for participants as well as model simulations. We ran 500 simulations of each hierarchical model (top) and flat model (bottom). See Table 1 for model parameters. Behavioral results show patterns of positive and negative transfer predicted by hierarchical, but not flat RL models, in both stages.

#### 2.2.2. Second stage choices reveal option transfer

To strengthen our results, we further examined the specific errors that participants made as they can reveal the latent structure used to make decisions. To further disambiguate between the Option Model and the Task-Set Model, we categorized errors into meaningful choice types ([9]). We focused on the second stage choices for model comparison (Fig. 4), the part of the experiment designed so that temporally extended policies could have an impact on decision making.

**Figure 4:**
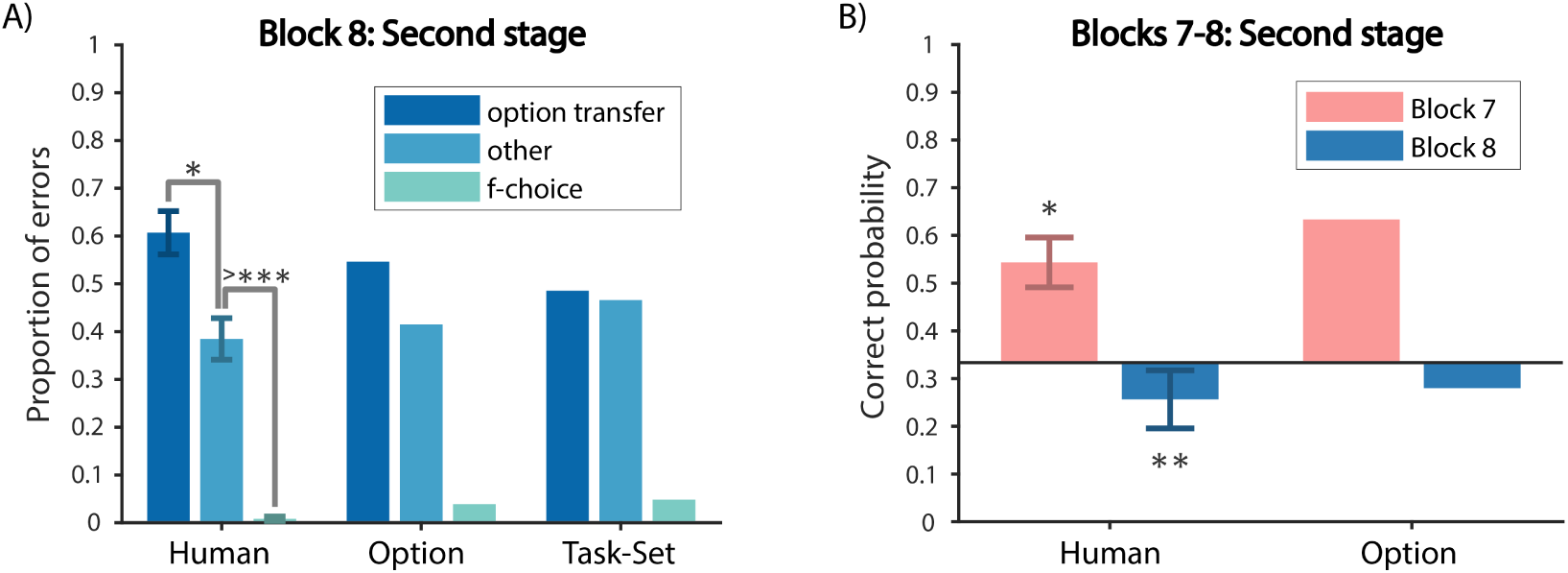
Experiment 1 second stage choices. (A) Error type analysis of the second stage in Block 8 for participants, the Option Model and the Task-Set Model. Participants made significantly more option transfer errors than other errors. This was predicted by the Option Model, but not by the Task-Set Model. (B) Probability of a correct first key press for the second stage of the first trial of each of the 4 branches in Blocks 7-8 reveals positive and negative transfer prior in first attempt (left), as predicted by the Option Model (right).

We hypothesized that participants learned *MO*’s that paired the policies in the first and second stages. Therefore, positive transfer in the second stage of Block 7 and negative transfer in the second stage of Block 8 should be due to participants selecting the entire *MO* that was previously learned in response to a first stage stimulus, including the correct key press for the first level stimulus as well as the corresponding *LO* for the second level. We defined choice types based on this hypothesis. For example, for the second stage of Block 8, consider the diamond following the circle in Block 8 (Fig. 1B): *A*_2_ is the correct action; an *A*_1_ error corresponds to the correct action in the first stage (“f-choice” type); an *A*_4_ error would be the correct action if selecting *MO*_1_ as a whole (“option transfer” type); an *A*_3_ error is labeled “other” type.

We computed the proportion of the 3 error types for the first 3 trials of each of the 4 branches in the second stage of Block 8 (Fig. 4A). There was a main effect of error type (1-way repeated measure ANOVA, *F*(2, 48) = 44, *p* < 0.0001). In particular, we found more “option transfer” errors than the “other” errors (paired t-test, *t*(24) = 2.5, *p* = 0.02), suggesting that participants selected previously learned *MO*’s as a whole at the beginning of the second stage of Block 8. The Option Model could reproduce this effect because the agent selects an entire option (*MO*) in the first stage: not only its immediate response to the first stage stimulus, but also its policy over *LO* choice in the second stage. The Task-Set Model could not reproduce this effect, because the first stage choice was limited to the first stage, and the second stage did not use any choice information from the first stage. Therefore, the error type profile in Block 8 could not be accounted for by transfer of one-step task-sets alone, ruling out the Task-Set Model.

There was also more “other” type than “f-choice” errors (paired t-test, *t*(24) = 8.8, *p* < 0.0001). There were few “f-choice” errors, likely due to meta-learning ([69]): participants observed that the correct action in the second stage was always different from the first stage (Fig. 1B). We included a mechanism in all models to capture this heuristic and quantitatively capture behavior better.

The same choice type definitions were not well-defined for the second stage of blocks other than Block 8. Therefore, we categorized errors differently in Blocks 1-7. For example, consider the diamond following the circle in Blocks 1, 3, and 5 (Fig. 1B): *A*_4_ is the “correct” choice; an *A*_1_ error corresponds to the correct choice in the first stage (“f-choice” type); an *A*_2_ error corresponds to the correct action for the other second stage stimulus, triangle, in the same *LO*, thus we defined it to be the “sequence” type, because *A*_2_ followed the first stage correct action *A*_1_ half of the time, as opposed to the “non-sequence” action *A*_3_, which never happened after *A*_1_. Aggregating the first 3 trials for each of the 4 branches in the second stage of Blocks 5-7 (Supplementary Fig. S6A), we did not find any significant difference in any of the 4 choice types between the second stage of Block 7 and that of Blocks 5-6 (paired t-test, all (*t*(24) ≤ 1, *p*’s> 0.30). This indicates that the positive transfer in the second stage of Block 7 was not interfered by the negative transfer in the first stage of Block 7, further confirming that participants were selecting learned *MO*’s as a whole, but re-composing them together into a new *HO*. The Option Model is also able to quantitatively capture the similarity of the choice type profiles between Block 7 and Blocks 5-6 (Supplementary Fig. S6B).

#### 2.2.3. The first press in the second stage reveals theoretical benefit of options

While the first several trials demonstrated transfer effects, the Option Model predicts immediate transfer effect on the first press in the second stage of a new block without any experience. Therefore, we computed the probability of a correct choice on the first press for the 4 branches in the second stage (Fig. 4B), and compared to chance (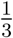, accounting for the meta-learning effect that the correct action in the second stage was always different from the first stage). The probability of a correct first key press in Block 7 and Blocks 5-6 was significantly above chance (sign test, Block 7: *p* = 0.015; Blocks 5-6: *p* < 0.0001), without significant difference between the two (sign test, *p* = 0.26). These positive transfer effects on the first press supports our prediction that participants were using previously learned *MO* to guide exploration and thus speed up learning even without any experience in Blocks 5-7. Block 8 was significantly below chance (sign test, *p* = 0.004), independently indicating, via negative transfer, exploration with previously learned *MO* in the very first trials. The Option Model was able to quantitatively reproduce these positive and negative transfer effects evident in the first press in the second stage, since the first stage choice can immediately help inform which *LO* to use in the second stage.

#### 2.2.4. First stage choices reveal transfer of policies over options

To test whether participants learned *HO*’s in the first stage, we investigated errors in the first stage. We hypothesized that the increase in key presses in the first stage of Block 7 (Fig. 2B) was due to selecting a previously learned but now wrong *HO* in the first stage, which would be characterized by a specific error. We categorized first stage errors into 3 types (“wrong shape”, “wrong *HO*”, and “both wrong”), which we exemplify for the circle in Blocks 1, 3, and 5 (Fig. 1B): *A*_1_ is the “correct” action; an *A*_2_ error corresponds to the correct action for the square in the same block (“wrong shape” type); an *A*_3_ error corresponds to the correct action for the circle in Blocks 2, 4, and 6 (“wrong *HO*” type); and *A*_4_ is the “both wrong” type. According to our hypothesis, we expected that the worse performance in the first stage of Block 7 (Fig. 3B) should be primarily due to the “wrong HO” errors. We found a main effect of choice type (2-way repeated measure ANOVA, *F*(3, 72) = 195, *p* < 0.0001) and a significant interaction between block and choice type (*F*(3, 72) = 2.9, *p* = 0.04). In particular, we found that in Block 7 (Fig. 5A), compared to Blocks 5-6, only the “wrong *HO*” error type marginally increased (paired t-test, *t*(24) = 1.9, *p* = 0.07) in Block 7. The Option Model reproduced this choice type profile in the first stage(Fig. 5B), by attempting to transfer previously learned *HO*, which would hurt performance in the first stage.

**Figure 5:**
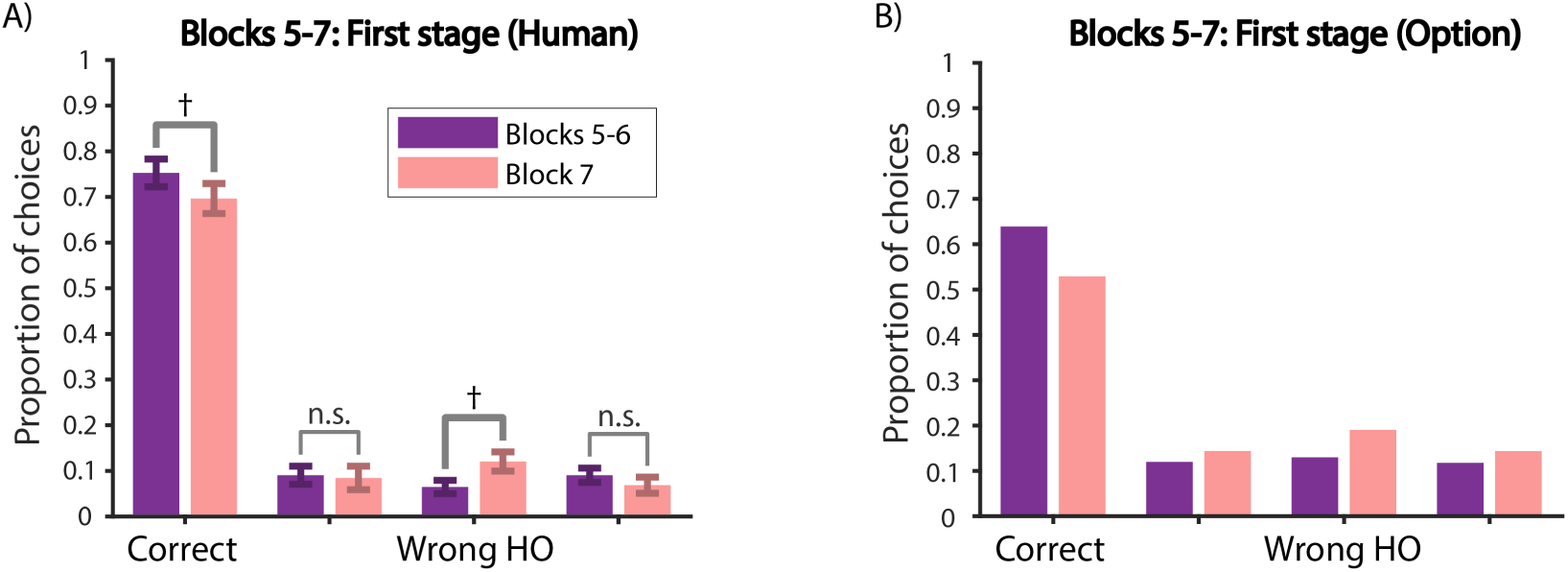
Experiment 1 first stage choices. Choice type analysis of the first stage in Blocks 5-7 for participants (A) and the Option Model (B). Participants made significantly more wrong *HO* errors in Block 7 than in Blocks 5-6, but no change for the other two error types. This suggests that participants were negatively transferring *HO* in the first stage of Block 7, as predicted by the Option Model.

#### 2.2.5. Experiment 1 Mturk replicates option transfer in the second stage

While in-lab participants’ behavior showed promising evidence in favor of transferring multi-step options, we sought to replicate our results in a larger and more diverse population. Therefore, we ran a shorter version of Experiment 1 on Mturk (Fig. 6A, Supplementary Fig. S12). In the second stage, we replicated the main effect of block on the number of presses (1-way repeated measure ANOVA, *F*(2, 108) = 19, *p* < 0.0001). Specifically, the average number of key presses (Fig. 6C) in the first 10 trials of Block 7 was not significantly different from that of Blocks 5-6 (paired t-test, *t*(54) = 0.72, *p* = 0.47). Participants pressed significantly more times in Block 8 compared to Block 7 and Blocks 5-6 (paired t-test, Block 7: *t*(54) = 4.5, *p* < 0.0001; Blocks 5-6: *t*(54) = 5.3, *p* < 0.0001), replicating results from in-lab participants (Fig. 2B).

**Figure 6:**
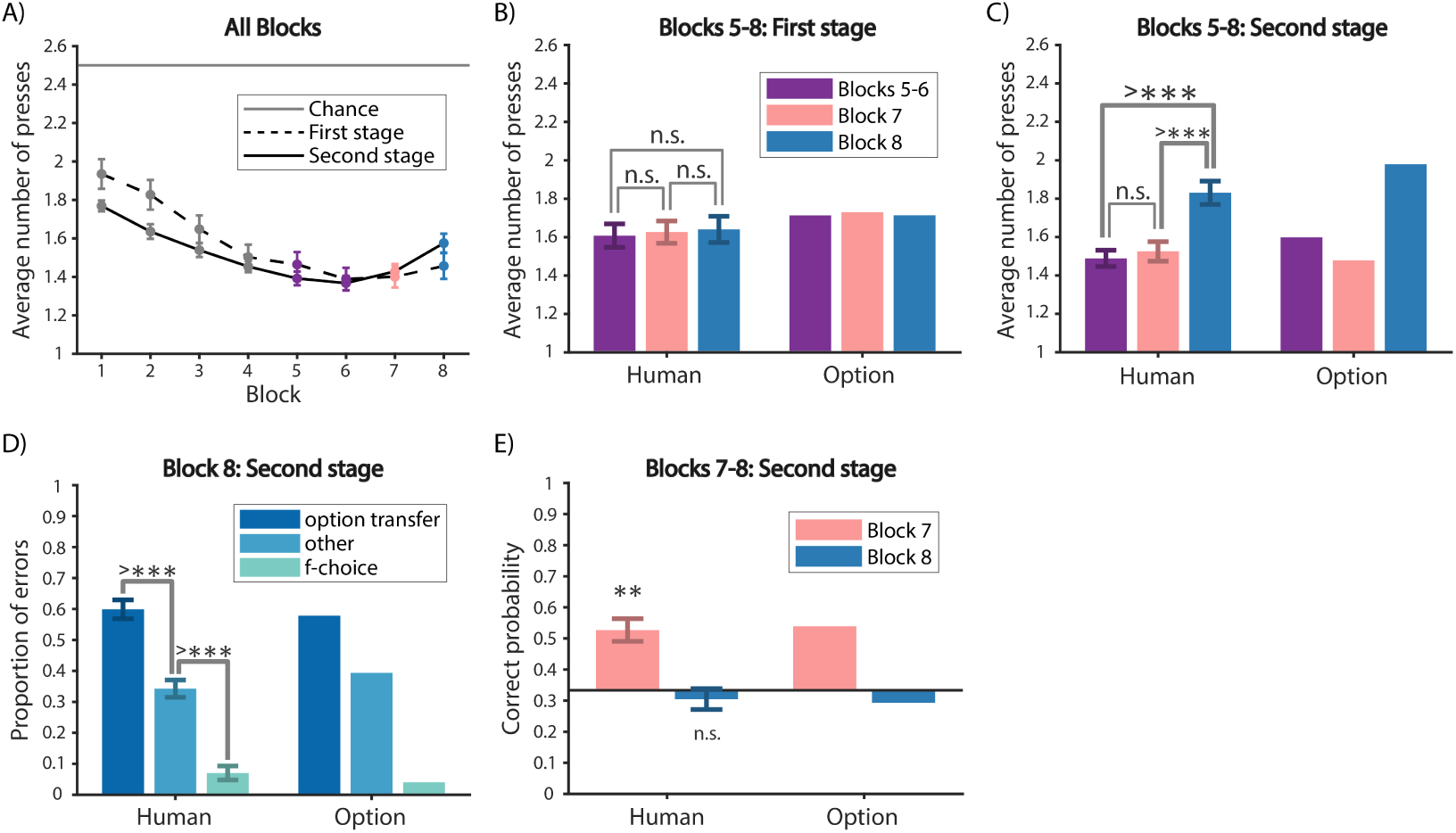
Experiment 1 Mturk results. (A) Average number of key presses in the first and the second stages per block. (B) Average number of key presses for the first 10 trials of Blocks 5-8 for the first stage for participants (left) and the Option Model (right). (C) Same as (B) for the second stage. (D) Error type analysis of the second stage in Block 8 for participants (left) and the Option Model (right). We replicated the same pattern as the in-lab population (Fig. 4A). (E) Probability of a correct first key press for the second stage of the first trial of each of the 4 branches in Blocks 7-8 for participants (left) and the Option Model (right).

In the second stage of Block 8 (Fig. 6D), there was a main effect of error type (1-way repeated measure ANOVA, *F*(2, 108) = 62, *p* < 0.0001). The “option transfer” errors were significantly more frequent than the “other” type errors (paired t-test, *t*(54) = 4.7, *p* < 0.0001), and the “other” type was significantly more frequent than the “f-choice” type (paired t-test, *t*(54) = 6.7, *p* < 0.0001). This also replicates the error type profile of in-lab participants.

For the probability of correct choice in the first press (Fig. 6E), we also found participants were performing significantly above chance in the second stage of Blocks 3-4, Blocks 5-6 and Block 7 (sign test, Blocks 3-4: *p* = 0.001; Blocks 5-6: *p* = 0.003; Block 7: *p* = 0.001), but not significantly different from chance in Block 8 (sign test, *p* = 0.18). There was also no significant difference between Block 7 and Blocks 5-6 (sign test, *p* = 1). This supported the previous finding that participants used temporally extended *MO*s to explore in a new context.

We did not replicate the negative transfer in the first stage of Block 7 (Fig. 6B) shown in in-lab participants (Fig. 2B). There was no main effect of block on the number of presses (1-way repeated measure ANOVA, *F*(2, 108) = 0.19, *p* = 0.83). Mturk participants did not press significantly more times in the first stage of Block 7 than Block 8 or Blocks 5-6 (paired t-test, Block 7: *t*(54) = 0.30, *p* = 0.77; Blocks 5-6: *t*(54) = 0.32, *p* = 0.75). This is potentially due to the lack of motivation among Mturk participants to exploit structure in the first stage, since participants did not receive points for being correct in the first stage. On the other hand, participants received points for choices in the second stage, which, as indicated by the Mturk experiment instruction, would impact their bonus. This might explain why the transfer effects in the first stage did not replicate, but the second stage transfer did. Note that in this case, the absence of transfer allowed the Mturk participants to make fewer errors in Block 7 than they might otherwise, highlighting the fact that engaging in a cognitive task and building and using structure is not always beneficial.

The option model was able to account for Experiment 1 Mturk data, despite the lack of transfer in the first stage, by assuming either a faster forgetting of *HO*s (higher *f* ^1^) or a lower prior for reusing them (higher *γ*^1^) (Table 1). Indeed, simulations reproduced the lack of transfer in the first stage (Fig. 6B), and also captured all option transfer effects demonstrated by Mturk participants in the second stage(Fig. 6C-E).

We conclude that, in the Mturk sample, similar to the in-lab sample, we successfully replicated the main option transfer effects in the second stage due to selecting a temporally extended policy *MO* as a whole. This is reflected by number of presses, proportion of error types in Block 8, and the probability of correct choice in the first press (Fig. 6C-E). While we did not replicate transfer of high level-options (task-sets of options), this could be accommodated by the model, and understood as a lack of motivation at learning the highest level of hierarchy *HO*.

## 3. Experiment 2

Experiment 2 was administered to UC Berkeley undergraduates in exchange for course credit. 31 (21 females; age: mean = 20.2, sd = 1.8, min = 18.3, max = 26.3) UC Berkeley undergraduates participated in Experiment 2. 4 participants in Experiment 2 were excluded due to incomplete data or below chance performance, resulting in 26 participants for data analysis.

### 3.1. Experiment 2 Protocol

Experiment 1’s Block 8 comes after a first testing block that includes re-composing of previous options, which could interfere with our interpretation of positive and negative transfer results in Block 8, for example by making participants aware of the potential for structure transfer. In Experiment 2, we removed Block 7 of Experiment 1 to eliminate this potential interference (Fig. 7). Therefore, Block 7 in Experiment 2 was identical to Block 8 in Experiment 1. In addition, to limit experiment length and loss of motivation at asymptote in each block, we decreased the length of Blocks 3-7 to 32 trials each, with each first stage stimulus leading to each second stage stimulus 8 times. All other aspects were identical to Experiment 1.

**Figure 7:**
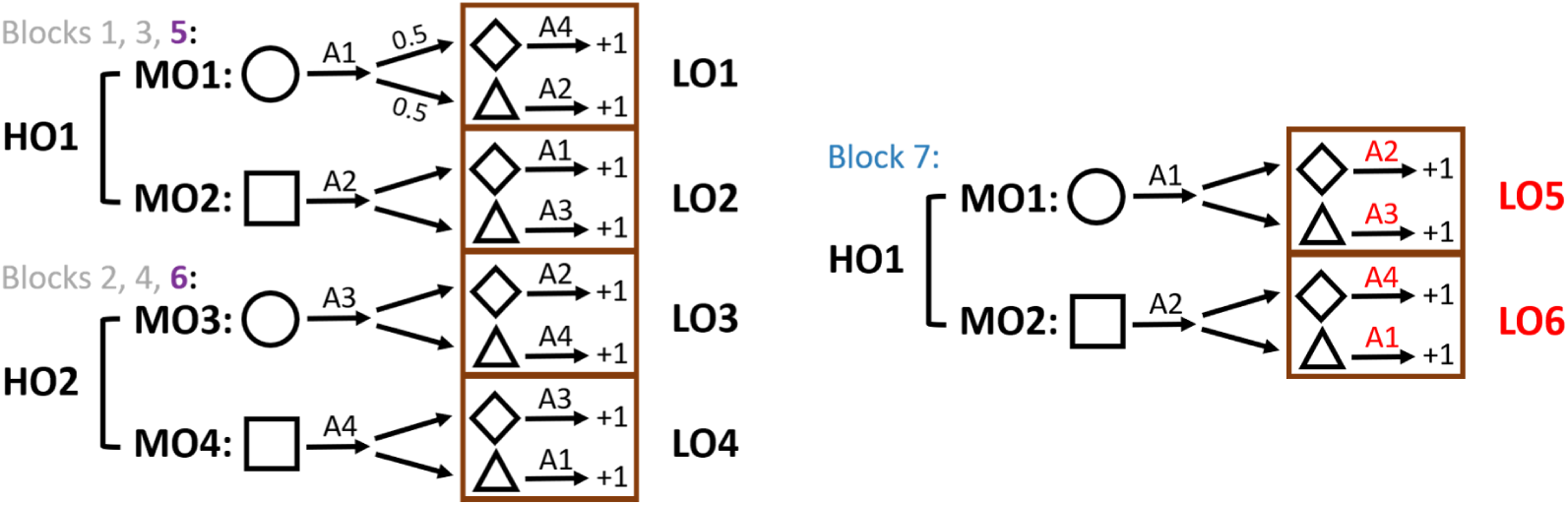
Experiment 2 protocol. To eliminate potential interference of Block 7 on Block 8 in Experiment 1, Block 7 of Experiment 1 was removed in Experiment 2. Therefore, Block 7 in Experiment 2 was identical to Block 8 in Experiment 1.

### 3.2. Experiment 2 Results

#### 3.2.1. Second stage choices replicate option transfer

Participants were able to learn the correct actions in both the first and second stages and their performance improved over Blocks 1-6, (Fig. 8A). The within-block learning curves also showed that participants performance improved and then reached asymptote as they progressed within a block (Supplementary Fig. S13).

**Figure 8:**
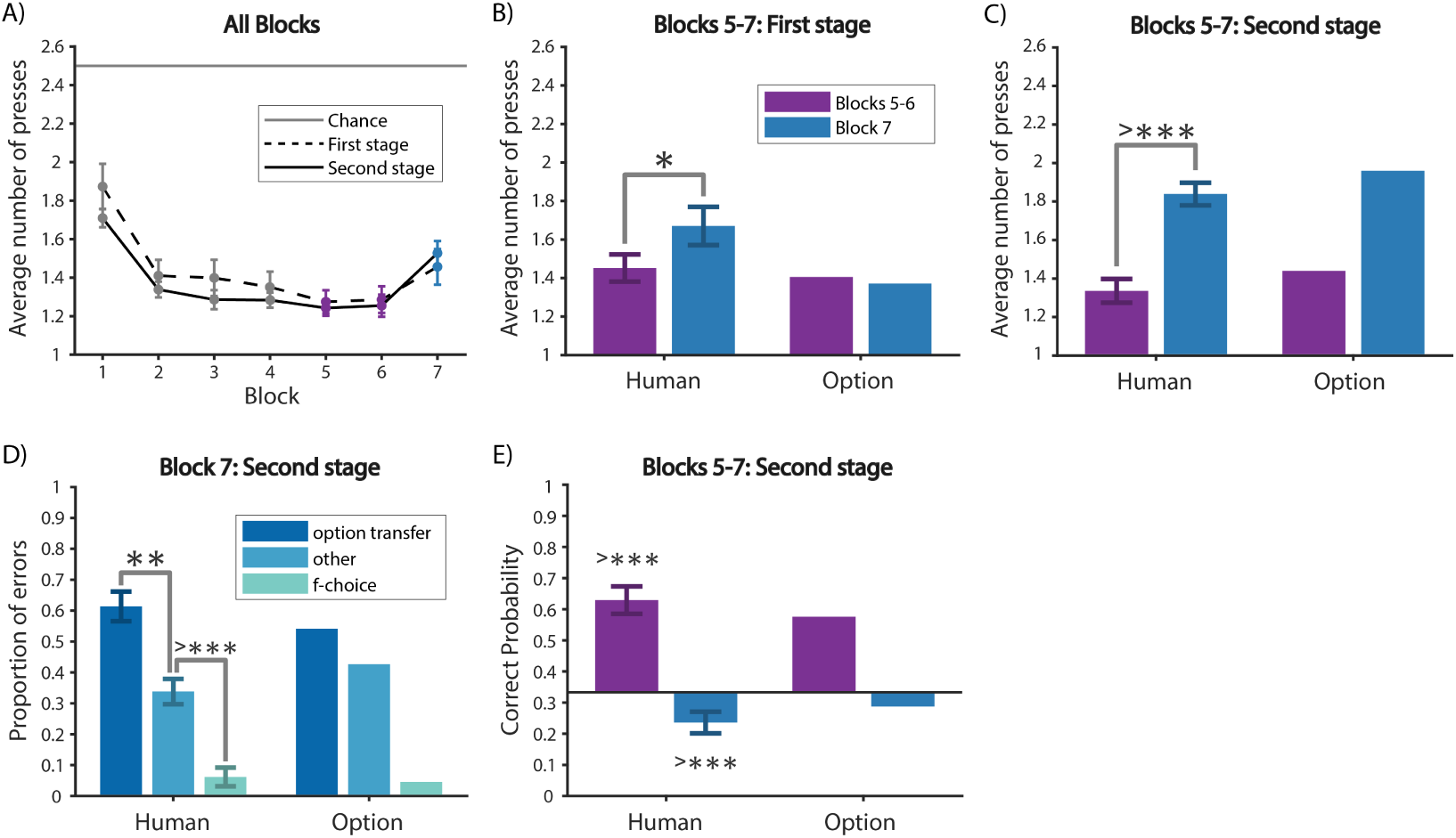
Experiment 2 results. (A) Average number of key presses in the first and the second stages per block. (B, C) Average number of key presses for the first 10 trials of Blocks 5-7 for the first (B) and second (C) stage for participants (left) and the Option Model (right). (D) Error type analysis of the second stage in Block 7 for participants (left) and the Option Model (right). We replicated the same pattern as in Block 8 of Experiment 1 (Fig. 4A, Fig. 6D). (E) Probability of a correct first key press for the second stage of the first trial of each of the 4 branches in Blocks 5-7 for participants (left) and the Option Model (right).

We replicated the negative transfer effects in the second stage of Experiment 1 (Fig. 2B) both in terms of number of presses (Fig. 8C) and error types in Block 7 (Fig. 8D). Participants pressed significantly more times in the second stage of Block 7 compared to Blocks 5-6 (paired t-test, *t*(25) = 6.4, *p* < 0.0001). In Block 7 specifically, there was a main effect of error type (1-way repeated measure ANOVA, *F*(2, 50) = 30, *p* < 0.0001). The proportion of the error type “option transfer” was significantly higher than the error type “other” (paired t-test, *t*(25) = 3.2, *p* = 0.004).

We also observed transfer effects on the first press in the second stage (Fig. 8E). We found that the probability of a correct choice was significantly above chance in Blocks 3-4 and Blocks 5-6 (sign test, Blocks 3-4: *p* = 0.0094; Blocs 5-6: *p* < 0.0001), and significantly below chance in Block 7 (sign test, *p* < 0.0001). This replicates results in Blocks 3-6 and 8 in Experiment 1 (Fig. 4B). The Option Model could quantitatively reproduce all these transfer effects (Fig. 8B-D).

#### 3.2.2. Second stage choices in Block 7 reveal interaction between meta-learning and option transfer

Because there was no Block 7 from Experiment 1, we had a less interfered test of negative transfer in the second stage of Block 7 of Experiment 2. Therefore, we further broke down the second stage choice types for each of the 4 branches in the second stage of Block 7 in Experiment 2 (Fig. 9A). Consider (Fig. 1B) the two first stage stimuli as *F*_1_ (circle) and *F*_2_ (square), and the two second stage stimuli as *S*_1_ (diamond) and *S*_2_ (triangle). We found a main effect of error type on proportion of error types and a marginally significant interaction between branch and error type (2-way repeated measure ANOVA, error type: *F*(2, 36) = 20, *p* < 0.0001; interaction: *F*(6, 108) = 2.1, *p* = 0.055). Specifically, we found the error type profile in Fig. 8C was mainly contributed by *F*_1_ → *S*_1_, i.e. circle in the first stage followed by diamond in the second stage, and *F*_2_ → *S*_2_ (paired t-test, *F*_1_ → *S*_1_: *t*(23) = 2.7, *p* = 0.013; *F*_2_ → *S*_2_: *t*(23) = 3.1, *p* = 0.005). On the other hand, there was no significant difference between the “option transfer” and “other” error types for *F*_1_ → *S*_2_ and *F*_2_ → *S*_1_ (paired t-test, *F*_1_ → *S*_2_: *t*(22) = 0.9, *p* = 0.38; *F*_2_ → *S*_1_: *t*(22) = 0.81, *p* = 0.43). It is striking that this highly non-intuitive result is perfectly predicted by the Option Model (Fig. 9B).

**Figure 9:**
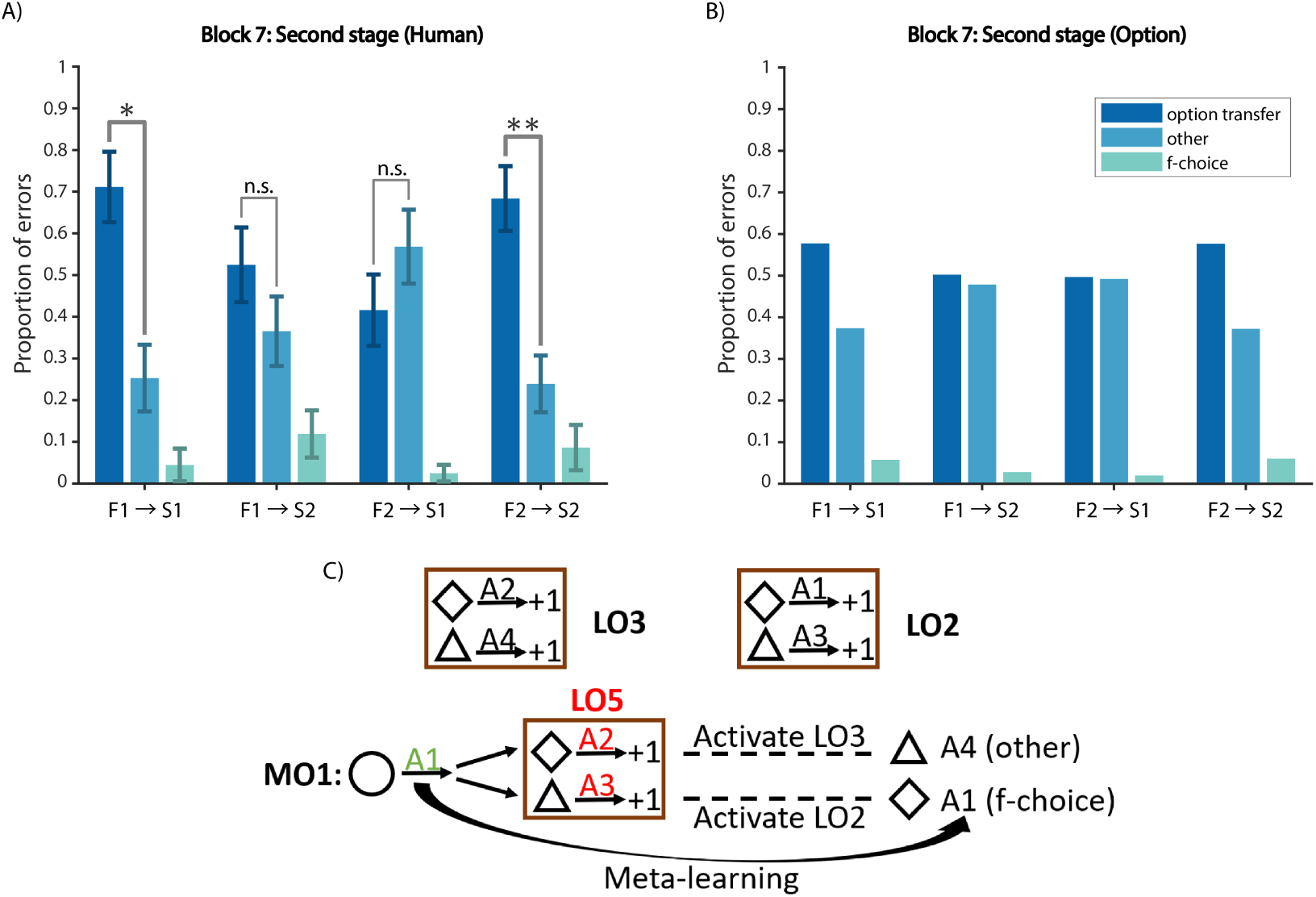
Experiment 2 second stage choice shows interaction between option transfer and meta learning. Error type analysis for each of the 4 branches in the second stage of Block 8 for participants (A) and the Option Model (B). The option transfer error was more than other error only for *F*_1_ → *S*_1_ and *F*_2_ → *S*_2_, which was predicted by the Option Model. (C) Example schematic for the interaction: learning *A*_2_ for the diamond activates *LO*_3_; learning *A*_3_ for the triangle activates *LO*_2_; meta-learning only suppresses *LO*_2_ but not *LO*_3_.

The Option Model offers an explanation as the interaction between option transfer and meta-learning (Fig. 9C). Meta-learning discourages participants from selecting second-stage actions that repeat the correct first-stage action, and as such, discourage them from sampling some, but not other *LO*s (e.g. *LO*_2_ in the example of Fig. 9C). This interference in the exploration of potential *LO*’s leads to some transfer errors to be more likely, in an asymmetrical way.

#### 3.2.3. Influence of the second stage on the first stage

For the first stage choices (Fig. 8B), we found that participants pressed significantly more times in the first 10 trials of Block 7 compared to Blocks 5-6 (paired t-test, *t*(25) = 2.4, *p* = 0.024). This effect was not found in Experiment 1 between Block 8 and Blocks 5-6 (Fig. 2B), and was not predicted by the model.

One potential explanation for this surprising result is that the error signals in the second stage propagated back to the first stage. Specifically, the errors participants made by selecting the wrong *LO* in the second stage are credited to the chosen *LO*’s policy, but participants might also credit these errors to using the wrong *HO* in the first stage. Going back to our example, if your meal is not tasty, it might not be because you roasted the potatoes instead of boiling them, but it might be because you needed vegetables instead of potatoes in the first place. To test this explanation, we further probed choice types in the first stage of Experiment 2 (Supplementary Fig. S7). Indeed, we found significantly more “wrong *HO*” errors in Block 7, compared to Blocks 5-6 (paired t-test, *p* = 0.045). Therefore, the increase in number of key presses in the first stage of Block 7 was mainly contributed by more “wrong *HO*” errors, indicating that participants explored another high level option (cooking vegetables). The same effect was not seen in the first stage of Experiment 1 between Block 8 and Blocks 5-6 (Fig. 2B), potentially due to the interference of Block 7 in Experiment 1.

The Option Model could not capture this effect, since the selection of *HO* was only affected by learning in the first stage (Sec. 2.1.5), as a way of simplifying credit assignment (see Sec. 6 for a more detailed discussion on credit assignment).

## 4. Experiment 3

Experiment 3 was administered to UC Berkeley undergraduates in exchange for course credit. 35 (22 females; age: mean = 20.5, sd = 2.5, min = 18, max = 30) UC Berkeley undergraduates participated in Experiment 3. 10 participants in Experiment 3 were excluded due to incomplete data or below chance performance, resulting in 25 participants for data analysis.

An additional 65 (37 female; see age range distribution in Table 3) Mturk participants finished the experiment. 34 participants were further excluded due to poor performance, resulting in 31 participants for data analysis (see Sec. 2.1.4).

### 4.1. Experiment 3 in-lab Protocol

In Experiment 1, to perform well in the second stage, participants had to learn option-specific policies, due to the non-Markovian nature of the task (the correct action for the same second stage stimulus was dependent on the first stage stimulus). In Experiment 3, we removed this non-Markovian feature of the protocol and tested whether the removal would reduce or eliminate option transfer. Based on previous research on task-sets showing that participants build structure when it is not needed ([32, 70]), we predicted that participants might still show some evidence of transfer. However, we predicted that any evidence of transfer would be weaker than in previous experiments.

In Experiment 3, the second stage stimuli following the two first stage stimuli were different (Fig. 10). For example, diamond and triangle followed circle, whereas star and hexagon followed square. This eliminated the key non-Markovian feature from Experiment 1, since participants could simply learn the correct key for each of the 4 second stage stimuli individually without learning option-specific policies. Blocks 1 and 2 had 60 trials; we shortened Blocks 3 to 8 to 32 trials for the same reason as in Experiment 2. All other aspects of the protocol were identical to Experiment 1.

**Figure 10:**
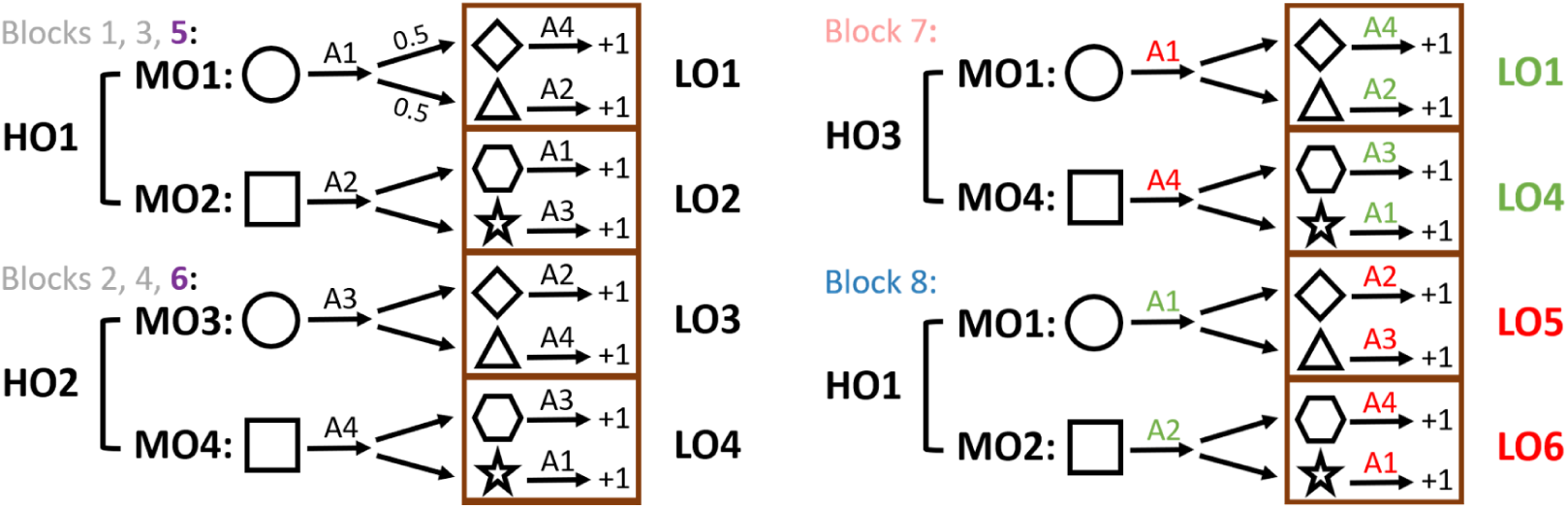
Experiment 3 protocol. The second stage stimuli following each first stage stimuli were different: diamond and triangle followed circle; hexagon and star followed square. All state-action assignments remained the same as Experiment 1. This manipulation allowed us to test whether participants would naturally learn and transfer options in the second stage even when they could simply learn the correct key for each of the 4 second stage stimuli individually, rather than needing to take into account first stage information.

### 4.2. Experiment 3 Mturk Protocol

In the Mturk version, Blocks 1 and 2 had a minimum of 32 and a maximum of 60 trials, but participants moved on to the next block as soon as they reached a criterion of less than 1.5 key presses per second stage trial in the last 10 trials (the 31 Mturk participants included for data analysis on average used 36 (SD = 7, median = 32, min = 32, max = 60) trials in Block 1 and 35 (SD = 4, median = 32, min = 32, max = 59) trials in Block 2). Blocks 3 to 8 all had 32 trials each. Experiment 3 MTurk was thus perfectly comparable to Experiment 1 MTurk, as such, we focus first on MTurk results, since the same comparison could not be drawn between Experiments 1 and 3 for in-lab participants.

### 4.3. Experiment 3 Results

#### 4.3.1. Mturk participants show reduced option transfer

Mturk participants were able to learn the correct actions in both the first and second stages, and their performance improved over Blocks 1-6, (Fig. 11A). The within-block learning curves also showed that participants performance improved and then reached asymptote as they progressed within a block (Supplementary Fig. S14).

**Figure 11:**
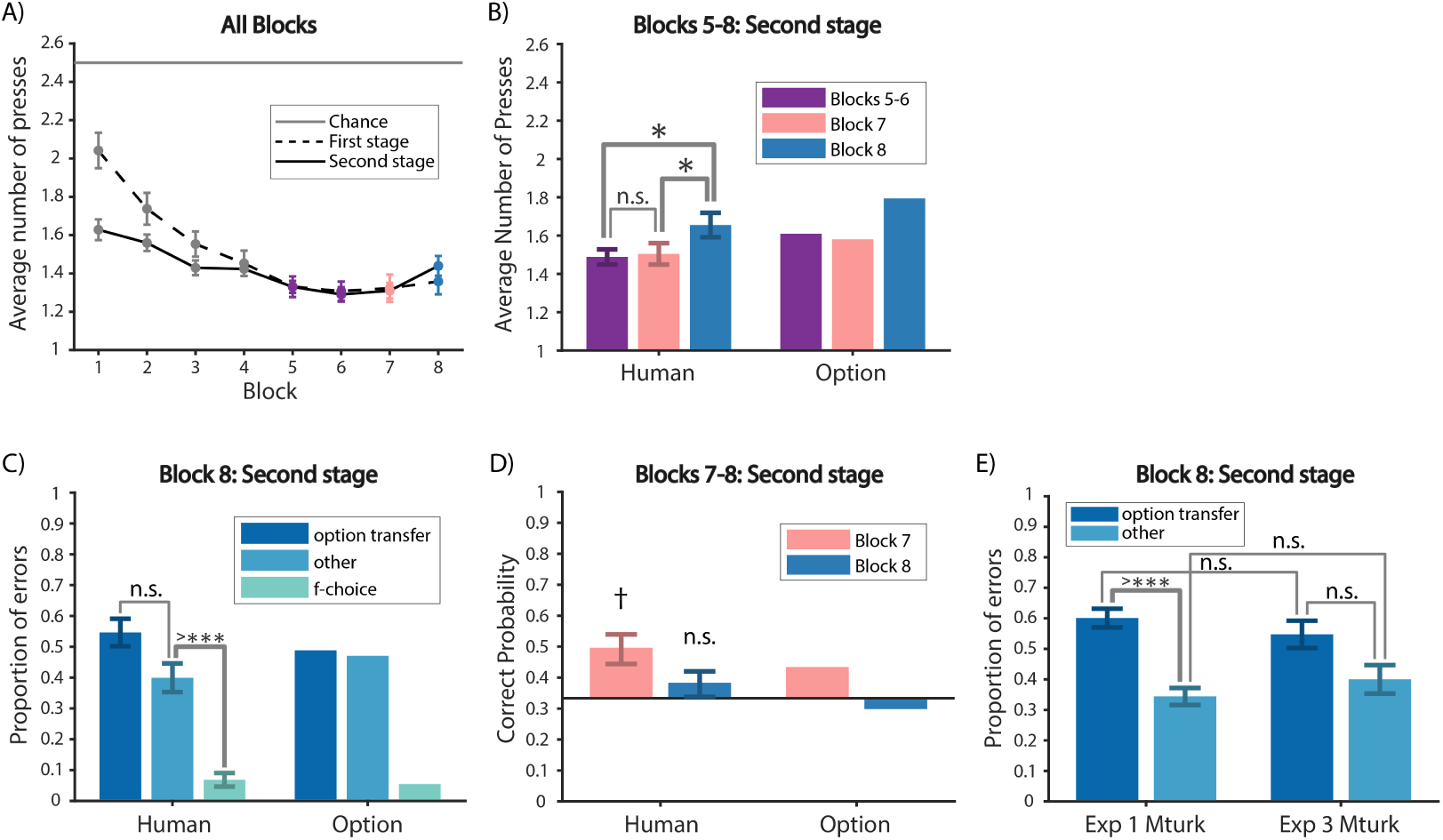
Experiment 3 Mturk results. (A) Average number of key presses in the first and the second stages per block. (B) Average number of key presses for the first 10 trials of Blocks 5-8 for the second stage for participants (left) and the Option Model (right). (C) Error type analysis of the second stage in Block 8 for participants (left) and the Option Model (right). The proportion of option transfer error was not significantly different from other error, different from Experiment 1 and Experiment 2, suggesting reduced option transfer. (D) Probability of a correct first key press for the second stage of the first trial of each of the 4 branches in Blocks 7-8 for participants (left) and the Option Model (right). (E) Comparison of Experiment 1 Mturk and Experiment 3 Mturk participants in terms of error types in the second stage of Block 8: There was no significant effect of experimental condition.

We first analyzed the average number of key presses in the first 10 trials of each block and stage. For the first stage (Supplementary Fig. S8A), we found no effect of block on number of presses across Blocks 5-8 (*F*(2, 60) = 0.13, *p* = 0.88), as in Experiment 1 MTurk. For the critical second stage (Fig. 11B), there was a main effect of Block (*F*(2, 60) = 3.3, *p* = 0.043). Specifically, there was no significant difference between Block 7 and Blocks 5-6 (paired t-test, *t*(30) = 0.25, *p* = 0.81). Participants pressed significantly more times in Block 8 than in Block 7 and Blocks 5-6 (paired t-test, Block 7: *t*(30) = 2.1, *p* = 0.048; Blocks 5-6: *t*(30) = 2.2, *p* = 0.036).

The negative transfer effect observed in the first stage of Block 7 in Experiment 1 (Fig. 3A) was not present here in Experiment 3 (Fig. 11). In addition to the fact that the first stage was never explicitly rewarded, as in Experiment 1, participants in Experiment 3 were even less motivated to exploit structure in the first stage. This is because the first stage in Experiment 3 was not necessary for resolving the second stage actions (Fig. 10), while the non-Markovian aspect of Experiment 1 (Fig. 1B) forced participants to incorporate first stage information to resolve the correct choice for the second stage.

We calculated the proportion of error types in the second stage of Block 8 (Fig. 11C). Unlike in Experiment 1, we did not observe significantly more “option transfer” error than “other” error (paired t-test, *t*(30) = 1.6, *p* = 0.11). This choice type profile, compared to that in Experiment 1 and Experiment 2 (Fig. 4A, Fig. 6D, Fig. 8D) suggests reduced option transfer in the second stage.

We also calculated the probability of a correct second stage first press for each of the 4 branches in the second stage (Fig. 11D). The probability was significantly above chance in Blocks 3-4 and Blocks 5-6 (sign test, Blocks 3-4: *p* = 0.0002; Blocks 5-6: *p* < 0.0001). It was marginally above chance in Block 7 (sign test, *p* = 0.07) and not significantly different from chance in Block 8 (sign test, *p* = 1). Compared to the results in Experiment 1 (Fig. 4B, Fig. 6E). These results suggest participants were still taking advantage of previously learned options to speed up learning at the beginning of each block, but potentially to a lesser extent compared to Experiment 1 and Experiment 2.

To formally quantify the effect of the experimental manipulation, we compared Experiment 1 and Experiment 3 for Mturk participants. In particular, we compared the proportion of “option transfer” and “other” error types in the second stage of Block 8 between the two experiments (Fig. 11E). We found a main effect of error type (2-way mixed ANOVA, *F*(2, 168) = 76, *p* < 0.0001), but there was no interaction between experiment and error type (2-way mixed ANOVA, *F*(2, 168) = 0.89, *p* = 0.41). In particular, the proportion of “option transfer” error type was not significantly higher in Experiment 1, compared to that in Experiment 3 (unpaired t-test, *t*(84) = 1, *p* = 0.32). This further shows that while there might be reduced option transfer in the second stage of Block 8 based on the error type profile (Fig. 11C), we could not rule out option transfer in Experiment 3.

The Option Model could capture a reduction in option transfer (Fig. 11B-D), with an increase in the second stage clustering coefficient *γ*^2^, which controls how likely the model is to select a new blank policy compared to previously learned ones in the second stage, as well as the forgetting parameter in the second stage, *f* ^2^, which increases the speed at which the model forgets previously learned *LO* (Table 1.

#### 4.3.2. In-lab participants replicate results from Mturk participants

In-lab participants replicated all aforementioned trends shown in Mturk participants (Supplementary Fig. S9). In particular, there was a main effect of block on number of choices in the second stage (*F*(2, 46) = 7.2, *p* = 0.002). In-lab participants also pressed significantly more times in the second stage of Block 8 than Blocks 5-6 (paired t-test, *t*(23) = 3.6, *p* = 0.0017), and marginally more than Block 7 (paired t-test, *t*(23) = 1.9, *p* = 0.067). Moreover, similar to Mturk participants, the proportion of “option transfer” error type was not significantly different from “other” error type (paired t-test, *t*(23) = 0.8, *p* = 0.43). These results replicated reduced option transfer in the second stage in a separate in-lab population. Note that we could not do the same comparison between Experiment 1 and Experiment 3 for in-lab participants, because the number of trials per block for Experiment 1 and Experiment 3 was different in-lab.

## 5. Experiment 4

Experiment 4 was administered to UC Berkeley undergraduates in exchange for course credit. 31 (23 females; age: mean = 20.2, sd = 1.4, min = 18, max = 23) UC Berkeley undergraduates participated in Experiment 4. 12 participants were excluded due to incomplete data or below chance performance, resulting in 19 participants for data analysis.

An additional 110 (50 females; see age range distribution in Table 3) Mturk participants finished the experiment. 49 participants were excluded due to poor performance, resulting in 61 participants for data analysis (see Sec. 2.1.4).

### 5.1. Experiment 4 in-lab Protocol

Experiment 4 (Fig. 12) was designed to test whether participants were able to compose options learned at different levels. Specifically, the protocol was identical to Experiment 1, except for Blocks 7 and 8. Block 8 in Experiment 4 was similar to Block 8 in Experiment 1, introducing two new *LO*’s (*LO*_*new*_) at the second stage as a benchmark for pure negative transfer.

**Figure 12:**
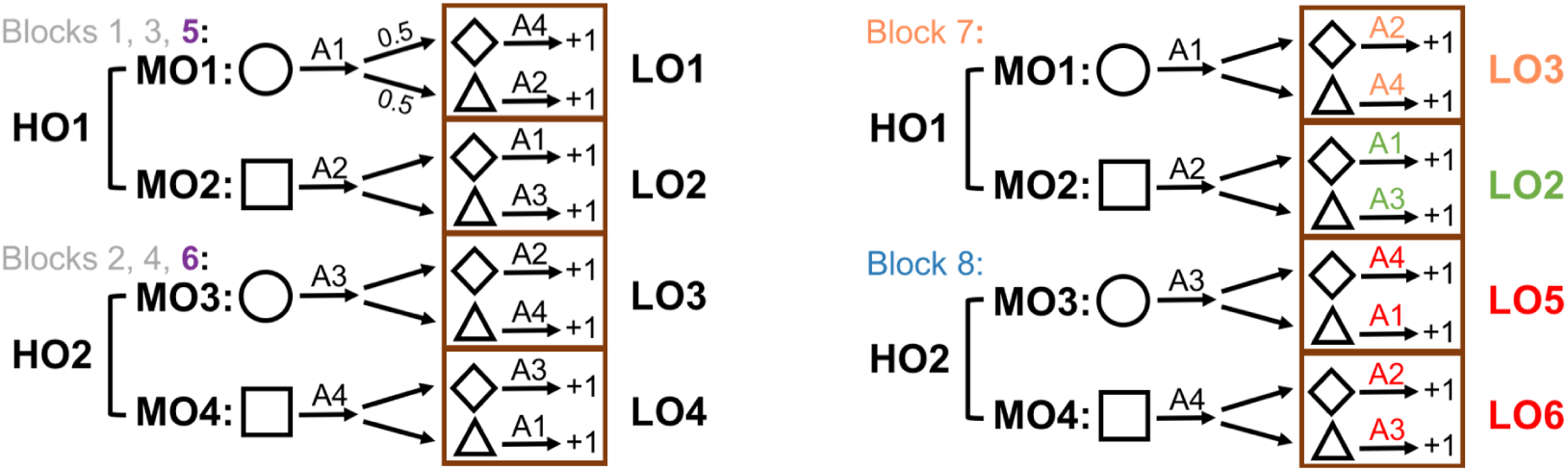
Experiment 4 protocol. In Experiment 4, we tested participants’ ability to recompose *LO* policies within *MO* policies. Blocks 1-6 were identical to Experiment 1. In Block 7, green indicates positions of potential positive transfer: *MO*_2_ followed by *LO*_2_ was learned in Blocks 1, 3, 5. Orange indicates positions of option composition: although *MO*_1_ previously included *LO*_1_ for second stage stimuli, it was modified to *LO*_3_ in Block 7. In Block 8, red indicates positions of negative transfer: *LO*_5_ and *LO*_6_ were completely novel to participants. Blocks were color coded for later analysis: Blocks 1-4 gray; Blocks 5-6 purple; Block 7 orange; Block 8 blue.

The main difference between Experiment 4 and Experiment 1 was Block 7. In Block 7, one of the first stage stimuli (e.g. square) elicited the same extended policy *MO*_2_ (*A*_2_ followed by *LO*_2_ in the second stage), allowing positive *MO* transfer (“match” condition *LO*_*match*_). In contrast, the other first stage stimulus (e.g. circle) elicited a new policy recomposed of old subpolicies: participants needed to combine what they learned in the first stage of *MO*_1_ in Blocks 1, 3, and 5 (*A*_1_) (allowing for first stage transfer of *HO*_1_), and the second stage of Blocks 2, 4, and 6 (*LO*_3_; “mismatch” condition *LO*_*mismatch*_). Extending the food analogy, in Blocks 1, 3, 5, participants learned to make potatoes (*MO*_1_) by cutting potatoes (the first stage) and then roasting (*LO*_1_). In Block 7, participants also needed to cut potatoes, but then steam them (*LO*_3_), which was already learned as part of *MO*_3_ (make vegetables) in Blocks 2, 4, 6. All blocks had 60 trials each.

#### 5.1.1. Experiment 4 Mturk Protocol

The Mturk version was shortened for online workers. Blocks 1 and 2 had a minimum of 32 and a maximum of 60 trials, but participants moved on to the next block as soon as they reached a criterion of less than 1.5 key presses per second stage trial in the last 10 trials (the 61 Mturk participants included for data analysis on average used 46 (SD = 11, median = 42, min = 32, max = 60) trials in Block 1 and 43 (SD = 11, median = 38, min = 32, max = 60) trials in Block 2). All other blocks had 32 trials each.

### 5.2. Experiment 4 Results

#### 5.2.1. Mismatch impacted performance of in-lab participants

Participants’ performance improved over Blocks 1-6 (Supplementary Fig. S10A) and within each block (Supplementary Fig. S16). First stage performance was similar in Blocks 5-8, as expected by the model (Supplementary Fig. S8). To test more specifically whether participants were able to compose options, we focused on comparing the second stage behavior for old *LO*s (*LO*_*match*_ and *LO*_*mismatch*_) and the average of *LO*_5_ and *LO*_6_ (*LO*_*new*_) in Blocks 7-8. The Option Model predicted that performance for *LO*_*match*_ in Block 7 should be the best due to positive transfer, since participants should have learned the extended *MO*_2_ policy whereby *LO*_2_ followed *A*_2_ in Blocks 1, 3, and 5 (Fig. 12). *LO*_*new*_ should be the worst due to negative transfer, with all 4 stimulus-action assignments in the second stage novel. Performance for *LO*_*mismatch*_ in Block 7 should fall in between (as observed in the number of key pressed, Fig. 13A). While there should be negative transfer, as *MO*_1_ was usually followed by *LO*_1_, *LO*_3_ had been previously learned, so its performance should still surpass the performance in the second stage of Block 8, where *LO*_5_ and *LO*_6_ were completely novel to the participants. Therefore, we predicted *LO*_*match*_ > *LO*_*mismatch*_ > *LO*_*new*_ in terms of performance.

**Figure 13:**
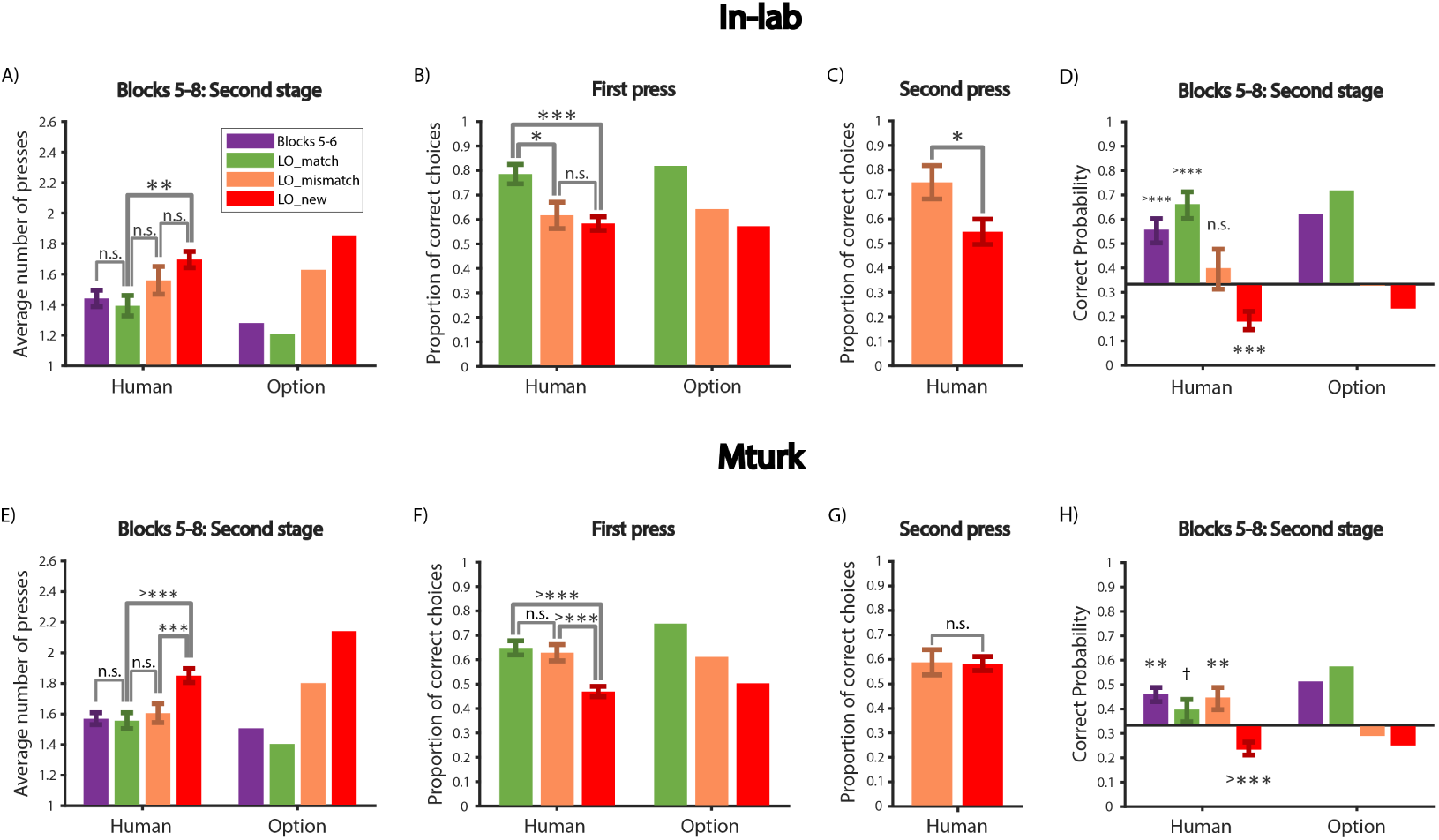
Experiment 4 results show re-composition of options. (A)-(D) In-lab participants. (A) Average number of key presses for the first 3 trials for each of the 4 branches in the second stage of Blocks 5-8 for participants (left) and the Option Model (right). Block 7 was split into *LO*_*match*_ and *LO*_*mismatch*_; Block 8 corresponded to *LO*_*new*_. (B) Proportion of correct choices on the first press of trials 1-3 for each of the 4 branches in the second stage for *LO*_*match*_, *LO*_*mismatch*_ and *LO*_*new*_ for participants (left) and the Option Model (right). (C) Proportion of correct choices on the second press (for trials 1-3 for each of the 4 branches with an incorrect first key press) for the mismatch (left) and the new (right) condition. (D) Probability of a correct first key press for the second stage of the first trial of each of the 4 branches in Blocks 5-8 for participants (left) and the Option Model (right). (E)-(H) Same as (A)-(D) for Mturk participants.

In the second stage (Fig. 13A), there was a main effect of block on number of presses (1-way repeated measure ANOVA, *F*(2, 36) = 9.9, *p* = 0.0004). Specifically, the average number of key presses in *LO*_*new*_ (Block 8) was significantly more than Blocks 5-6 and *LO*_*match*_ (paired t-test, Blocks 5-6: *t*(18) = 4.1, *p* = 0.0007; *LO*_*match*_: *t*(18) = 3.6, *p* = 0.002). There was no significant difference between Blocks 5-6 and *LO*_*match*_ (paired t-test, *t*(18) = 0.7, *p* = 0.49), supporting the model’s prediction of positive *MO* transfer in this condition. The model predicted that *LO*_*mismatch*_ performance should be between *LO*_*new*_ and *LO*_*match*_: *LO*_*mismatch*_ performance should reflect positive *LO* transfer but negative *MO* transfer. This was observed qualitatively, though the results did not reach significance (paired t-test, *LO*_*match*_: *t*(18) = 1.6, *p* = 0.13; *LO*_*n*_*ew*: *t*(18) = 1.4, *p* = 0.18). These results replicate the negative transfer effects in the second stage of Block 8 shown in Experiment 1 (Fig. 4A) and Experiment 2 (Fig. 8D). In addition, they provide initial support for the compositionality hypothesis of the model, with intermediary transfer in the mismatch condition.

We confirmed the previous results by analyzing the proportion of trials in which the first key press was correct. We found that, in the first 3 trials for each of the 4 branches in the second stage (Fig. 13B), there was a main effect of *LO* condition (1-way repeated measure ANOVA, *F*(2, 36) = 7.2, *p* = 0.002) on the proportion of correct choices for the first press of each trial. In particular, we found no significant difference between *LO*_*mismatch*_ and *LO*_*new*_ (paired t-test, *t*(18) = 0.56, *p* = 0.58), while the performance of *LO*_*match*_ was significantly higher than *LO*_*mismatch*_ and *LO*_*new*_ (paired t-test, *LO*_*mismatch*_: *t*(18) = 2.6, *p* = 0.017; *LO*_*new*_: *t*(18) = 4.4, *p* = 0.0003). These results suggested that the mismatch between *MO*_1_ and *LO*_3_ impacted participants’ performance, a marker of negative option (*MO*) transfer. In the first three iterations, participants’ first presses indicated that they were not able to efficiently re-compose the *LO*_*mismatch*_ into a new mid-level option.

To better investigate participants’ choices before they experienced any new information in a new block, we also computed the probability of a correct first key press for the second stage of the first trial of each of the 4 branches in the Blocks 5-8 (Fig. 13D). We found a main effect of block (Friedman Test, *χ*^2^(2, 36) = 20, *p* < 0.0001). Specifically, Blocks 5-6 and *LO*_*match*_ were significantly above chance (sign test, both *p* < 0.0001); *LO*_*mismatch*_ was not significantly different from chance (sign test, *p* = 0.34); *LO*_*new*_ was significantly below chance (sign test, *p* = 0.0007). There was a marginal difference between *LO*_*match*_ and *LO*_*mismatch*_ (sign test, *p* = 0.09), but no significant difference between *LO*_*mismatch*_ and *LO*_*new*_ (sign test, *p* = 0.24). These results further showed that the mismatch condition impacted participants’ performance on the first press due to negative option (*MO*) transfer, and replicated the strong negative transfer in Block 8 in Experiment 1 and Experiment 2. The Option Model captured participants’ behavior well (Fig. 13ABD, see Table 1 for model parameters).

#### 5.2.2. Second press reveals benefit of option composition

The results so far supported one of our predictions, *LO*_*match*_ > *LO*_*mismatch*_, by showing that performance in the mismatch condition was impacted due to negative *MO* transfer. We next sought evidence for our second prediction, *LO*_*mismatch*_ > *LO*_*new*_, where we hypothesized better performance in the mismatch condition by composing the first stage policy of *MO*_1_ and *LO*_3_.

In terms of performance on the first press in each trial, we did not found a significant difference between the two conditions (Fig. 13B). However, this might be because the negative *MO* transfer reduced the benefit of compositionality, making it less detectable on the first press, also reflected by the small effect from the Option Model in Fig. 13B. Positive *LO* transfer thus might only show a more significant effect after the first press unexpectedly failed (from negative transfer of *MO*_1_).

Therefore, we further computed the proportion of correct choices on the second press in those trials where the first press was incorrect (Fig. 13C). Indeed, we found that the proportion of correct choices on the second press was significantly higher in the mismatch condition than the new condition (paired t-test, *t*(17) = 2.8, *p* = 0.012). This result supports our second prediction, *LO*_*mismatch*_ > *LO*_*new*_, revealing a benefit in the mismatch condition compared to the new condition in participants re-composing an old *LO* into a non-matching *MO*.

#### 5.2.3. Mturk participants showed benefits of option composition

We collected a larger and independent sample on Mturk. Mturk participants also improved over Blocks 1-6 (Supplementary Fig. S10B) and within block (Supplementary Fig. S17), though their asymptotic performance (Blocks 5-6) was lower than the in-lab population. Specifically, we compared the average number of key presses in Blocks 5-6 in the first and second stages for both in-lab and Mturk populations. There was a main effect of stage and a marginal interaction of population and stage (2-way mixed ANOVA, stage: *F*(1, 78) = 7.1, *p* = 0.009; interaction: *F*(1, 78) = 3.1, *p* = 0.08). In particular, for the first stage, Mturk population was not significantly worse than the in-lab population (unpaired t-test, *t*(78) = 0.17, *p* = 0.86); but for the second stage, which was the focus of our analysis, Mturk population was significantly worse than the in-lab population (unpaired t-test, *t*(76) = 3.2, *p* = 0.002).

In the second stage (Fig. 13E), there was a main effect of block on number of presses (*F*(2, 120) = 17, *p* < 0.0001). Specifically, the average number of key presses in *LO*_*new*_ was significantly more than *LO*_*match*_ and *LO*_*mismatch*_ (paired t-test, *LO*_*match*_: *t*(60) = 4.6, *p* < 0.0001; *LO*_*mismatch*_: *t*(60) = 3.8, *p* = 0.0004). *LO*_*match*_ was not significantly different from Blocks 5-6 and *LO*_*mismatch*_ (paired t-test, Blocks 5-6: *t*(60) = 0.26, *p* = 0.8; *LO*_*mismatch*_: *t*(60) = 0.8, *p* = 0.42).

The proportion of correct first press choices (Fig. 13F) showed a similar pattern: there was a main effect of *LO* condition (*F*(2, 120) = 15, *p* < 0.0001) on the proportion of correct choices. In particular, the proportion of correct choice for *LO*_*new*_ was significantly lower than *LO*_*mismatch*_ and *LO*_*match*_ (paired t-test, *LO*_*mismatch*_: *t*(60) = 4.7, *p* < 0.0001; *LO*_*match*_: *t*(60) = 5.1, *p* < 0.0001) in Block 7. There was no significant difference between *LO*_*mismatch*_ and *LO*_*match*_ performance (paired t-test, *t*(60) = 0.54, *p* = 0.59). There was no difference between the mismatch condition and the new condition for second key presses (paired t-test, *t*(52) = 0.08, *p* = 0.94, Fig. 13G), contrary to in-lab participants (Fig. 13C). This difference could be attributed to MTurk participants’ lower task engagement. Indeed, contrary to in lab participants, MTurk participants’ performance was at chance for second key press (MTurk: paired t-test, *t*(53) = 1.6, *p* = 0.13; in-lab *t*(17) = 3.4, *p* = 0.003). Directly comparing MTurk and in-lab population for the proportion of correct second key press in both the mismatch and new conditions revealed a marginal effect of condition and a marginal interaction of population and condition (2-way mixed ANOVA, condition: *F*(1, 69) = 3.3, *p* = 0.07; interaction: *F*(1, 69) = 3.7, *p* = 0.06). This supports our interpretation that MTurk participants did not attempt to find the correct answer following an error, making the second press error analysis in this population difficult to interpret.

Finally, we looked at the probability of a correct first press in the very first trial of each of the 4 branches in the second stage (Fig. 13H). There was a main effect of block (Friedman test, *χ*^2^(2, 120) = 17, *p* = 0.0002). In particular, Blocks 5-6 and *LO*_*mismatch*_ were significantly above chance (sign test, both *p* = 0.004)l *LO*_*match*_ was marginally above chance (sign test, *p* = 0.07); *LO*_*new*_ was significantly below chance (sign test, *p* < 0.0001).

These results can be interpreted in one of two ways. The similar performance between *LO*_*match*_ and *LO*_*mismatch*_ suggests that participants were able to efficiently re-compose the first stage of *MO*_1_ with *LO*_3_ in the mismatch condition in Block 7, so that they did not suffer from *MO* negative transfer, as did in-lab participants. Alternatively, this result might indicate a lack of *MO* transfer (and only positive *LO* transfer) in both the match and mismatch condition. The latter interpretation is supported by the fact that second stage performance in *LO*_*match*_ was lower in MTurk participants than it was for in-lab participants in all measures (unpaired t-test, number of key presses in the first 10 trials of Blocks 5-6: *t*(78) = 1.8, *p* = 0.08; proportion of correct choices in match condition: *t*(78) = 2.4, *p* = 0.019).

The Option Model could capture the negative transfer effect in *LO*_*new*_ and thus the difference between *LO*_*new*_ and *LO*_*mismatch*_ (Fig. 13EF). However, it could not fully reproduce the lack of difference between *LO*_*match*_ and *LO*_*mismatch*_, since the model would first try to transfer *LO*_1_ in the mismatch condition, resulting in worse performance for *LO*_*mismatch*_.

This interpretation might suggest that the Task-Set Model explains the Mturk population better, indicating a lack of temporally extended options, and makes a specific prediction: second stage errors should not be impacted by first stage information. To test this prediction, we analyzed the specific errors participants made, as this is a specific hallmark of temporally extended option transfer vs. task-sets (Fig. 4A). Contrary to the prediction made by the Task-Set model, but consistent with the Option Model prediction, Mturk participants did demonstrate the behavioral signature of negative option (*MO*) transfer in the mismatch condition (Fig. 14): they made significantly more “option transfer” errors than “other” errors (paired t-test, *t*(53) = 4.8, *p* < 0.0001). While the comparison was not significant for in-lab participants (paired t-test, *t*(17) = 1.5, *p* = 0.16), a direct comparison between in-lab and Mturk populations did not reveal an effect of population (2-way mixed ANOVA, *F*(2, 140) = 0.74, *p* = 0.48). Thus, our results indicate that both MTurk participants and in-lab participants used temporally extended *MO*s, although MTurk participants were overall less successful at transferring them to facilitate decision making in the second stage. The results are consistent with participants re-composing low-level options into higher-level options.

**Figure 14:**
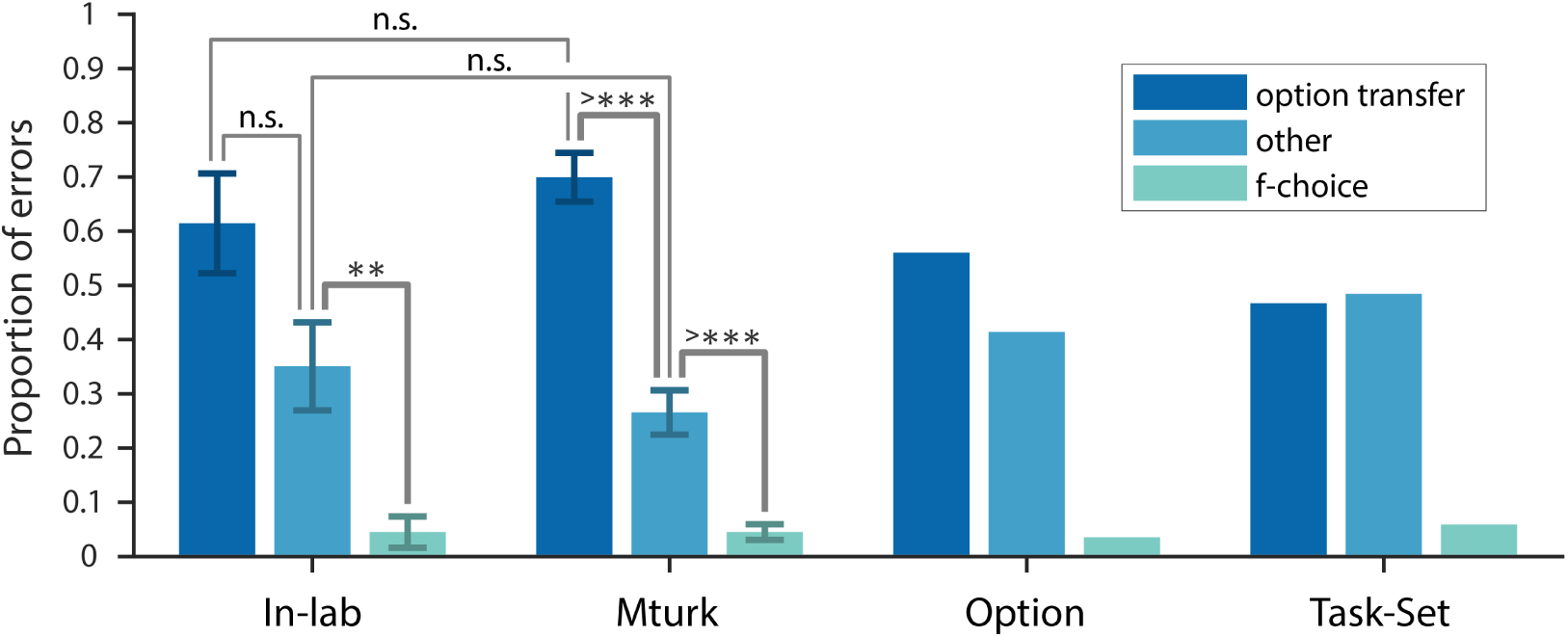
Experiment 4 second stage errors reveal temporal options transfer and compositionality. Error type analysis of the second stage in Block 7 for the mismatch condition for in-lab participants, Mturk participants, the Option Model and the Task-Set Model.

## 6. Discussion

Our findings provide novel and strong support for the acquisition of options in healthy human adults. Options can be thought of as choices that are more abstract than simple motor actions, but can be taken as a single choice. Using a novel two-stage protocol, we provide evidence that humans create options, and flexibly transfer and compose previously learned options. This transfer and composition ability guides exploration in novel contexts and speeds up learning when the options are appropriate, but impairs performance otherwise, as predicted by the options framework [11]. Model simulations showed that only a model including temporal hierarchy could account for all results, suggesting that human participants not only build state abstractions with one-step task-sets ([67]), but also temporal abstractions in the action space with multi-step options.

We developed a new model, the Option Model, to account for participants’ behavior. The Option Model includes features from our previous hierarchical structure learning model ([9, 32, 66]) and the hierarchical reinforcement learning (HRL) options framework ([43]). In our previous hierarchical structure learning model, we used non-parametric priors (CRP) over latent variables that represented the currently valid policy to create *state abstractions*: this allowed the model to cluster different contexts together if the same task-set applied. This CRP prior enables the agent to identify (via Bayesian inference) novel contexts as part of an existing cluster if the cluster-defined task-set proves successful, resulting in more efficient exploration and faster learning.

On the other hand, the original formulation of the HRL options framework ([43]) augments the action space of traditional flat RL with *temporal abstractions* called options. Each option is characterized by an initiation set that specifies which states the option can be activated, a termination function that maps states to a probability of terminating the current option, and an option-specific policy (that leads the agent to a potentially meaningful and useful subgoal).

Our Option Model is inspired by the fact that task-sets and options are similar in essentials: they are policies that an agent can select as a whole, and then apply at a lower level of abstraction (applying it to make a motor choice in response to a stimulus for task-sets, or applying it across time until termination in the case of an option [cite my structure learning book chapter]). Thus, our model brings together state and temporal abstractions by using option-specific CRP priors to implement option-specific policies that can be flexibly selected in different contexts if they share the same environmental contingencies. Our model captures the essence of the options framework despite some subtle differences. Here, we discuss how our Option Model relates to each part of the HRL options framework.

### Initiation set

The initiation set specifies the set of states where an option can be selected. The observable states in our tasks are the shapes shown on the screen. Therefore, at first, the initiation sets of *HO* and *MO* are first stage stimuli (e.g. circle and square, Fig. 1B), whereas the initiation sets of *LO* are second stage stimuli. However, the optimal policies were also dependent on the block; thus participants needed to infer the hidden context (*state abstraction*) dictated by block. Our CRP implementation can thus be thought of as continuously adding new block contexts to the initiation set of an option throughout the task. The ability to add new contexts to the initiation sets provides our Option Model the crucial flexibility needed to achieve transfer and composition, as demonstrated by human participants. For example, if *LO*_3_ was tied solely to the context of Block 2, where it was first learned, we would not observe the benefit of option composition in Experiment 4 in the mismatch condition.

### Termination function

An option’s termination function maps each state to the probability of terminating the current option (i.e. not using its policy anymore). How to terminate an option is closely related to the underlying theoretical question of credit assignment, which arises naturally in tasks that require hierarchical reasoning ([71]): if the current policy does not generate any (pseudo-) reward for a while, should the agent continue improving the current policy or terminate it and use another policy or even something new?

With a termination function as described in the original HRL options framework, credit assignment happens in a very specific way: the policy of the currently selected option (or options if multiple nested options are selected) is updated until termination is reached. In our task, this would make behavior very inflexible. For example, when an agent entered the second stage of Block 8 in Experiment 1 (Fig, 1B) for the first time after having correctly made a choice for the circle in the first stage, the agent would likely use *LO*_1_ due to negative transfer of *MO*_1_ and thus not receive reward. Because the termination function only takes state as an input, the agent would keep overwriting the *LO*_1_ policy with *LO*_5_ policy until termination, and thus not be able ot reuse *LO*_1_ down the line.

Our Option Model, however, uses a more flexible form of option termination. Specifically, we use Bayesian inference (Sec. 2.1.5), which was introduced in our previous hierarchical structure learning model ([9]). At the end of each choice, the model updates the likelihood of each option being valid based on the observed reward feedback, which then determines whether the model should stop using the current option. Moreover, Q-learning only operates on the option that has the highest posterior, thus assigning credit retrospectively to the best cause ([72]). Therefore, the Option Model is more likely to create a new *LO*_5_ and learn its policy from scratch, making it more flexible at learning and selecting options.

The crucial difference between the two is that the Option Model would create a new *LO*_5_ and learn its policy from scratch, without overwriting the original *LO*_1_ policy. While the Option Model can capture participants’ choices well across all four experiments, the current experimental protocol was not designed specifically to test credit assignment to options, and could not distinguish between these two possibilities. This remains an important question for future research.

There is another credit assignment problem that is not fully addressed by our current protocol and modeling: choices by lower level options may affect the termination of higher level options. For example, if you get punished for boiling potatoes, should you credit this to the lower level option (boiling) or to the higher level option (making potatoes in the first place). Should you plan to cook vegetables instead, or just roast the potatoes? We have some evidence for both levels of credit assignment (e.g. in Block 7 of Experiment 2, or Block 8 in Experiment 1; Fig. 1B), when participants were experiencing many errors in the second stage using *LO*_1_ and *LO*_2_. Participants might not only consider terminating or re-learning the current *LO*, but also naturally attribute some of the negative feedback to the choices they made in the first stage regarding *MO* or *HO*. Indeed, we observed that second stage errors potentially resulted in more “wrong *HO*” errors in the first stage of Experiment 2 (Supplementary Fig. S7).

In our Option Model (Sec. 2.1.5), for simplicity, first stage choices were only determined by learning within the first stage and were not sensitive to reward feedback in the second stage. It will be important in future research to better understand interactions between option levels for credit assignment. When considered together with the termination problem, these future directions may help trace the underlying neural mechanisms for credit assignment in human learning and hierarchical decision making.

### Option-specific policy

The most important component of an option is the option-specific policy: what lower level-choices (either simpler options or basic actions) it constrains. In this paper, we focused on the transfer of option-specific policy to test theoretical benefits of the options framework.

Theoretical work ([11]) suggested that useful options should facilitate exploration and speed up learning. Indeed, we observed speed up in learning through the positive transfer effects. For example, in Experiment 1, the second stage of Block 7 provided a test of positive option transfer in terms of both number of presses (Fig. 2B) and choice types (Supplementary Fig. S6). Importantly, this positive transfer was not interfered by the negative transfer in its first stage (Fig. 2B), suggesting that participants transferred mid-level options (*MO*) as a whole.

Moreover, the learning benefit was evident even in the first press (Fig. 4B, Fig. 6E, Fig. 8D): participants were already significantly above chance in the first press, indicated that they could explore by immediately transferring previously learned options.

Previously learned option-specific policies also helped with option composition in the mismatch condition (Fig. 12) of Experiment 4 (Fig. 13). While *MO*_1_ was usually followed by *LO*_1_ in Blocks 1, 3, 5, in the mismatch condition, *MO*_1_ was followed by *LO*_3_ instead. This change indeed resulted in “option transfer” errors (Fig. 14). However, the fact that *LO*_3_ had been previously learned helped participants explore more efficiently. For example, once participants figured out *A*_2_ was correct for the diamond, they would more likely explore *LO*_3_, and thus *A*_4_ for triangle.

The HRL options framework also suggested that non-useful options can slow down learning. Indeed, we observed negative option transfer effects in the second stage across multiple experiments in terms of number of presses (Fig. 2B, Fig. 6C, Fig. 8C, Fig. 13AE), and more importantly, error types (Fig. 4A, Fig. 6D, Fig. 8D, Fig. 9, Fig. 14), that are consistent with the predictions of the options framework. Note that the slow down was due to negative transfer of previously learned option-specific policies. Thus testing how having a wrong subgoal can impact learning performance is an interesting future direction.

We sought to confirm that participants were indeed learning option-specific policies, not just action sequences. Our protocol specifically used two second stage stimuli following each first stage stimulus (Fig. 1B) to avoid this potential confound. If, for example, circle was always followed by diamond and square by triangle, participants would not need to pay attention to the actual stimulus in the second stage, and could instead plan a sequence of actions in the first stage. In contrast, here, participants could only perform well by selecting options (i.e. stimulus-dependent temporally extended policies). While pure sequence learning could not account for our results, we investigated whether it could contribute to some of its aspects. Sequence learning would predict faster reaction times for actions that often follow in a sequence ([73]). Therefore, we compared the reaction time for the “sequence” and “non-sequence” error types in the second stage (Sec. 9.2). We did not find significant difference between the reaction time for “sequence” and “non-sequence” error types at the beginning of blocks; we only found such difference at the end of blocks (Supplementary Fig. S1, Fig. S2, Sec. 9.2). This suggests that while the transfer effects we observe at the beginning of each block could not be explained by pure sequence learning, participants might develop sequence learning-like expectations over time in a block, speeding up choices that came more frequently after each other.

We tested predictions of HRL options framework through positive and negative transfer of option-specific policies in the simplest possible set up of tabular representation of state and action space. Multiple aspects could be expanded on in future research to increase the generalizability of the policy in real world scenarios. First, real world policies apply to much more complex (continuous, multidimensional) state spaces. Recent work in AI expands the options framework to more realistic situations ([74]), where artificial agents learn how to navigate a sequence of rooms with different shapes and sizes. If each state in a room is naively paramatrized in a tabular way by (x, y) coordinates, when the agent is placed in a new room of a different shape, previously learned policy would be of not use. It is thus crucial to identify meaningful features of the state space shared by different rooms. ([74]) proposed learning options in a state space parametrized by distance from goals (“agent space”) to bypass this limitation.

Second, the low-level action space in real life conditions is also more complex. A good example is our flexible use of tools ([75]). We can conceptualize using various tools as taking actions. Humans demonstrate great flexibility when improvising using different tools to solve the same problem or even crafting new tools. If we simply represent actions in a tabular way, after participants associated a particular tool (action) to solve a task, the policy would be of no use if this particular tool is no longer provided in the future. The key might again be figuring out meaningful dimensions of the tool (action) space that are shared in different task scenarios, such as shape and weight of the tool.

Finally, even if two problems are different in terms of both state and action space (e.g. learning to play piano vs learning to play violin ([38])), knowledge of one might still help the other. Once one learned a piece on the piano, the knowledge of music theory might serve as a model to guide option transfer when learning the same piece on violin. These are important future directions for testing how humans transfer in those more real life scenarios, which might provide insight into developing more flexible and human-like AI systems with the HRL options framework.

### Option discovery

One of the most important questions regarding options in AI is how to discover meaningful options. Discovering useful options entails learning all components of an option: initiation set, termination function, and option-specific policy that leads to a meaningful sub-goal. In this paper, we designed a protocol that focused on learning option-specific policies by making all other features, including subgoals, trivial.

Discovering options may be useful because of a key feature of our interactions with our environment. In real world scenarios, it is frequent that for a given observable state, the right choice to make depends on hidden context, task demand, or past information. This property is refered to as *non-Markovian*: the current observable information is insufficient to determine the next step. For example, when potatoes are peeled, we can use them to make either roasted potatoes or mashed potatoes. Therefore, the state *“peeled potatoes”* is a meaningful subgoal state, and peeling potatoes is its corresponding option-specific policy.

This non-Markovian property might contribute to the hierarchical and compositional nature of human behavior. It is central to the original formulation of the options framework ([43]), and is also a natural objective for option discovery. In relation to our protocol, the correct action for diamond (Fig. 1B) varies from time to time in the same block. It makes sense to create different options to capture this, and relate it to the inferred hidden cause for why the correct actions change. Indeed, we observed that the non-Markovian feature in our experiments encouraged participants to create and transfer options at multiple levels of abstractions.

We tested whether the environment needs to be non-Markovian to trigger option creation. Specifically, we designed Experiment 3 by eliminating the non-Markovian property from Experiment 1 and testing if that affects option learning and transfer (Fig. 11). Unsurprisingly, we found weaker option transfer effects in Experiment 3; however, participants’ behavior was still not flat (Fig. 11, Supplementary Fig. S9). Thus, our results hint at the possibility that participants create temporal options (*MO*), even in the absence of a need for it, echoing past results showing that humans tend to create structure unnecessarily ([9, 70, 76, 77]). Furthermore, this may also show that objectives for option discovery are not limited to solving non-markovian problems. For example, ([12]) showed that humans could identify bottleneck states from transition statistics, reflecting graph-theoretic objectives for option discovery in humans.

### The options framework and other learning systems

While our Option Model uses a simple form of model-free RL (Q-learning; [1]) to learn option-specific policies, the options framework is general and not limited to just Q-learning. Options can be learned or used with model-free methods ([11]) and model-based methods ([44]). It also has strong connections to successor representations ([78, 79]), which might provide objectives for subgoal discovery.

Moreover, in this paper, we gave examples of potential interaction of options with the meta-learning system (Fig. 9) and sequence learning (Sec. 9.2) in human participants. How options might interact with other learning systems is an important question for future research.

## 7. Conclusion

In summary, we found compelling evidence of option learning and transfer in human participants by examining the learning dynamics of a novel two-stage experimental paradigm. Through analyzing participants’ behavioral patterns and model simulations, we demonstrated the flexibility of option transfer and composition at distinct levels in humans.

Humans’ ability to flexibly transfer previously learned skills is crucial for learning and adaptation in complex real world scenarios. This ability is also one of the fundamental gaps that sets humans apart from current state-of-the-art AI algorithms. Therefore, our work trying to probe learning and transfer in humans might also help provide inspirations for AI algorithms to be more flexible and human-like.

## 8. Acknowledgements

We thank Katya Brooun, Ham Huang, Helen Lu, Sarah Master, and Wendy Shi for their substantial contribution to the project. We thank Rich Ivry, Milena Rmus and Amy Zou for feedback on this draft. This work was supported by NIMH RO1MH119383.

## 9. Supplement

### 9.1. Potential asymmetry in Block 7 of Experiment 1

We checked whether the performance of circle and square in Block 7 was asymmetrically affected due to the interleaving of odd and even blocks (Fig. 1B). Specifically, participants might start Block 7 by using *HO*_1_ in odd blocks; thus the negative transfer in the first stage of Block 7 would be primarily due to more key presses from the square, not the circle.

To test this possibility, we calculated average number of key presses in the first 5 trials for circle and square respectively in Block 7. However, we found no significant difference between the performance of circle and square in the first stage (paired t-test, *t*(24) = 1.38, *p* = 0.18); we also found no significant difference between the performance in the second stage following circle and square (paired t-test, *t*(24) = 0.44, *p* = 0.66).

### 9.2. Second stage reaction time and sequence learning effects

Sequence learning ([73]) predicts that the reaction time of the “sequence” type to be faster than the “non-sequence” type. Therefore, we calculated the average reaction time (Fig. S1) for both “sequence” and “non-sequence” error types in Experiment 1 and 2.

#### 9.2.1. Experiment 1

**Supplementary Figure S1:**
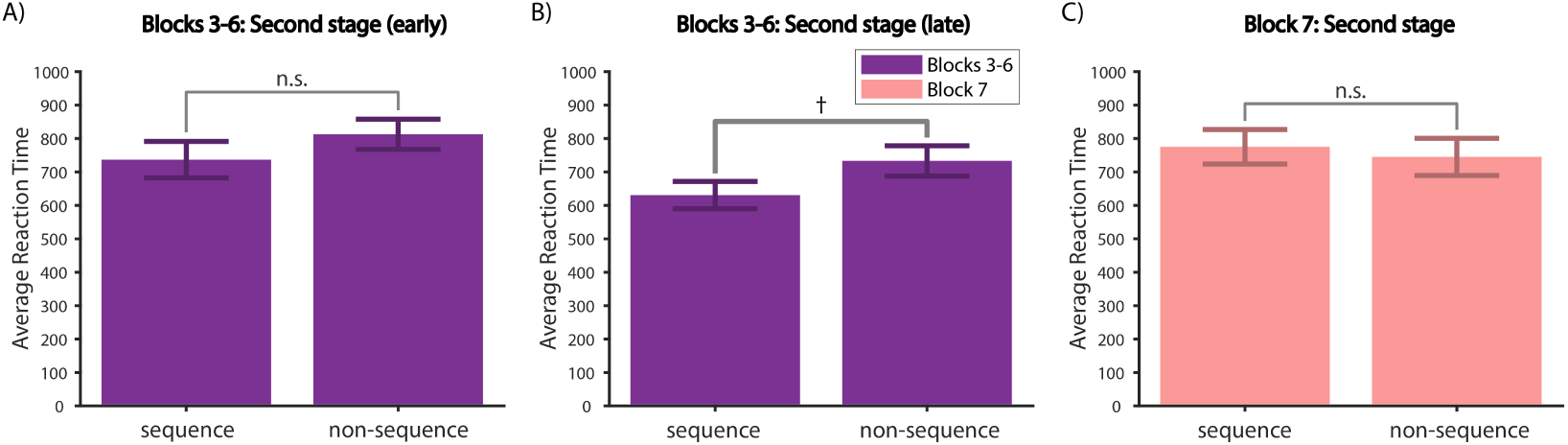
Experiment 1 reaction time. (A) Average reaction time for trials 1-7 for each of the 4 branches in the second stage for Blocks 3-6 for sequence (left) and non-sequence (right) error types. (B) Same as (A) for trials 8-15. (C) Average reaction time for sequence (left) and non-sequence (right) error types in the second stage of Block 7.

We broke down each block to 2 different time periods: early (trials 1-7 for each of the 4 branches in the second stage) and late (trials 8-15 for each of the 4 branches). Aggregating Blocks 3-6, we found a marginal effect of time period (2-way repeated measure ANOVA, *F*(1, 21) = 3.0, *p* = 0.099), which might be due to participants generally becoming faster as they progressed within a block. We also found a main effect of error type (2-way repeated measure ANOVA, *F*(1, 21) = 4.5, *p* = 0.046) on reaction time. Specifically, we found no significant difference (*t*(23) = 1.3, *p* = 0.2) between the reaction time of the “sequence” and “non-sequence” error types in the early time periods (Supplementary Fig. S1A). The “sequence” type was marginally faster (paired t-test, *t*(22) = 1.9, *p* = 0.072) than the “non-sequence” type in the late time period (Supplementary Fig. S1B). We also found no significant difference (paired t-test, *t*(20) = 1.1, *p* = 0.3) between the “sequence” and “non-sequence” types in the entire Block 7 (Supplementary Fig. S1C). These results suggest that the transfer effects we observed at the beginning of each block could not be due to pure sequence learning, which only start to take effect during learning saturation.

#### 9.2.2. Experiment 2

**Supplementary Figure S2:**
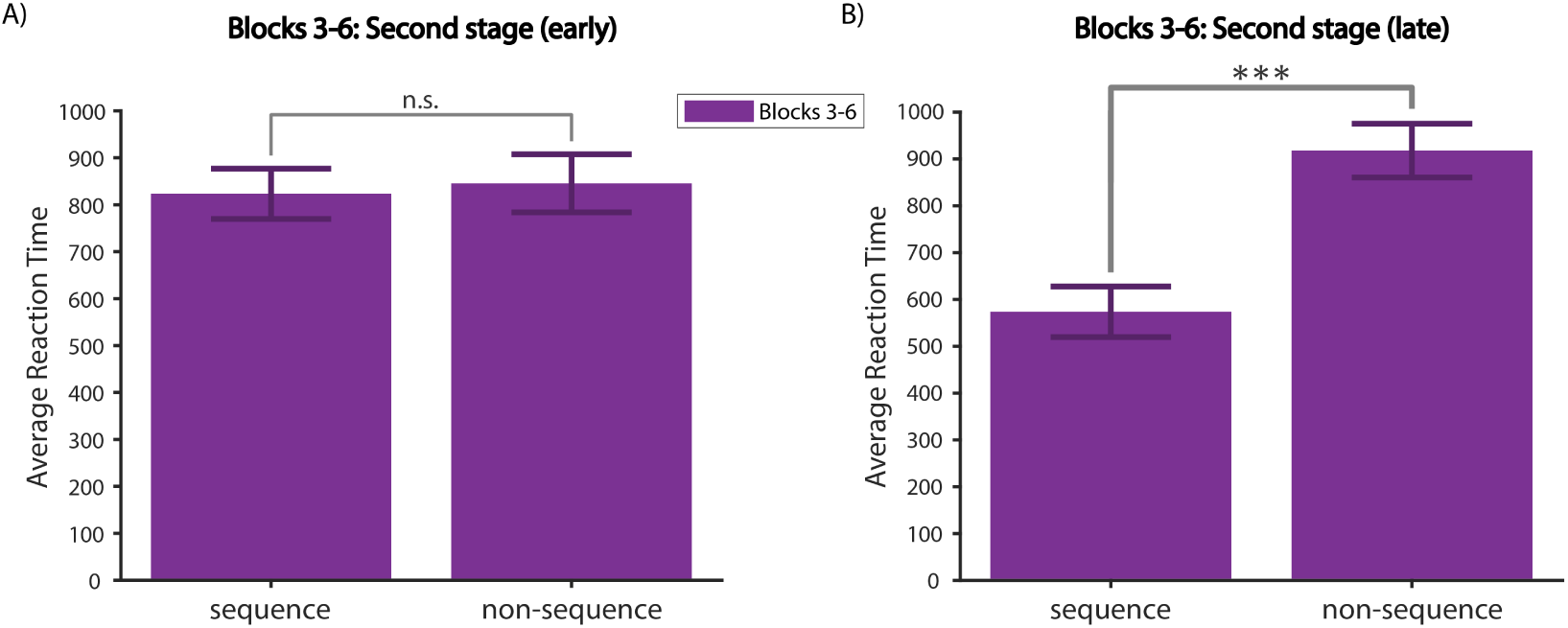
Experiment 2 reaction time. (A) Average reaction time for trials 1-4 for each of the 4 branches in the second stage for Blocks 3-6 for sequence (left) and non-sequence (right) error types. (B) Same as (A) for trials 5-8.

We also analyzed the reaction time (Fig. S2) of the “sequence” and “non-sequence” error types in Blocks 5-6 in Experiment 2. As in Experiment 1, we broke down each block into 2 halved time periods: early (trials 1-4 for each of the 4 branches in the second stage) and late (trials 5-8 for each of the 4 branches). We found a main effect of time period and error type, and a significant interaction (2-way repeated measure ANOVA, time period: *F*(1, 16) = 8, *p* = 0.012; error type: *F*(1, 16) = 16, *p* = 0.0009; interaction: *F*(1, 16) = 15, *p* = 0.0013). Specifically, there was no significant difference (Supplementary Fig. S2A) between the reaction time of the “sequence” and “non-sequence” types in the early time period (paired t-test, *t*(21) = 0.61, *p* = 0.55). However, the “sequence” type was significantly faster (Supplementary Figure S2B) than the “non-sequence” type in the late period (paired t-test, *t*(17) = 4.8, *p* = 0.0002). These results replicated the trend observed in the second stage of Experiment 1 (Supplementary Fig. S1A-B): sequence learning might take effect during learning saturation, but not the beginning of blocks, where we typically expect to observe transfer effects.

### 9.3. Parameters for model simulations

#### 9.3.1. Parameters used for main text

We used the set of parameters from Table 1 in the main text to track participants’ behavioral patterns both qualitatively and quantitatively.

#### 9.3.2. A set of constrained parameters that capture behavior across all tasks qualitatively

In the main text, we selected parameters to try to trace participants’ behavior patterns both quantitatively and qualitatively (Table 1). Here we used another set of parameters (Table 2) to (1) constrain parameters so that most experiments shared the same parameters while showing the qualitatively trends in participants’ behavior and (2) show that the model can reproduce the same qualitative effects with a range of parameters.

In particular, we used *α*^1^ = 0.7, *β*^1^ = 4, *β*^2^ = 4, *m* = 0.01 for all experiments. For all in-lab experiments, we used *α*^2^ = 0.7, *f* ^2^ = 0.001; for all Mturk experiments, we used *α*^2^ = 0.5, *f* ^2^ = 0.005, which indicate slower learning rate and faster forgetting. For Experiment 1 in-lab, we used *γ*^1^ = 14, *f* ^1^ = 0.001; for all other experiments, we used *γ*^1^ = 100, *f* ^1^ = 0.01 to implement a lack of transfer effects in the first stage. We used *γ*^2^ = 20 in Experiment 3 to model reduced option transfer in the second stage; for all other experiments, we used *γ*^2^ = 4.

We recreated some of the representative analysis in the main text to demonstrate that this second set of parameters can replicate the transfer effects in human participants qualitatively well.

**Supplementary Figure S3:**
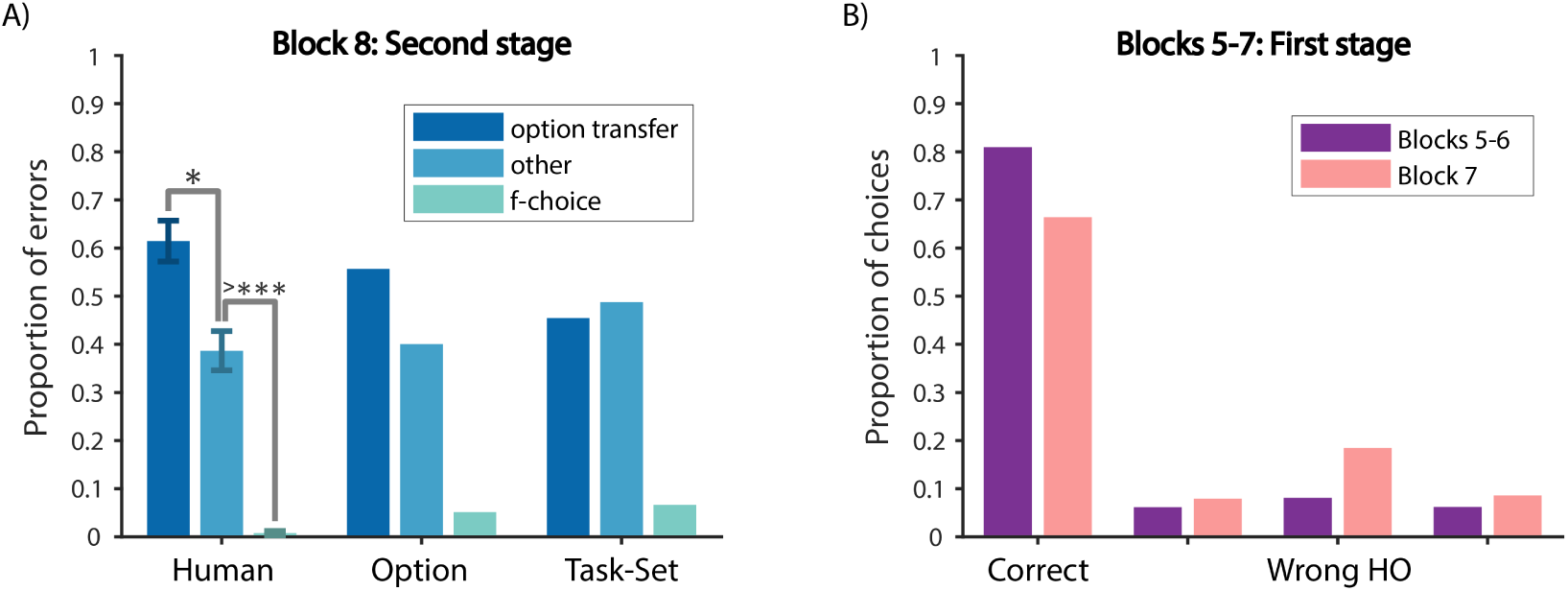
Experiment 1 with parameters from Table 2. (A) Error type analysis of the second stage in Block 8 for participants (left), the Option Model (middle) and the Task-Set Model (right). (B) Choice type analysis of the first stage in Blocks 5-7 for the Option Model.

**Supplementary Figure S4:**
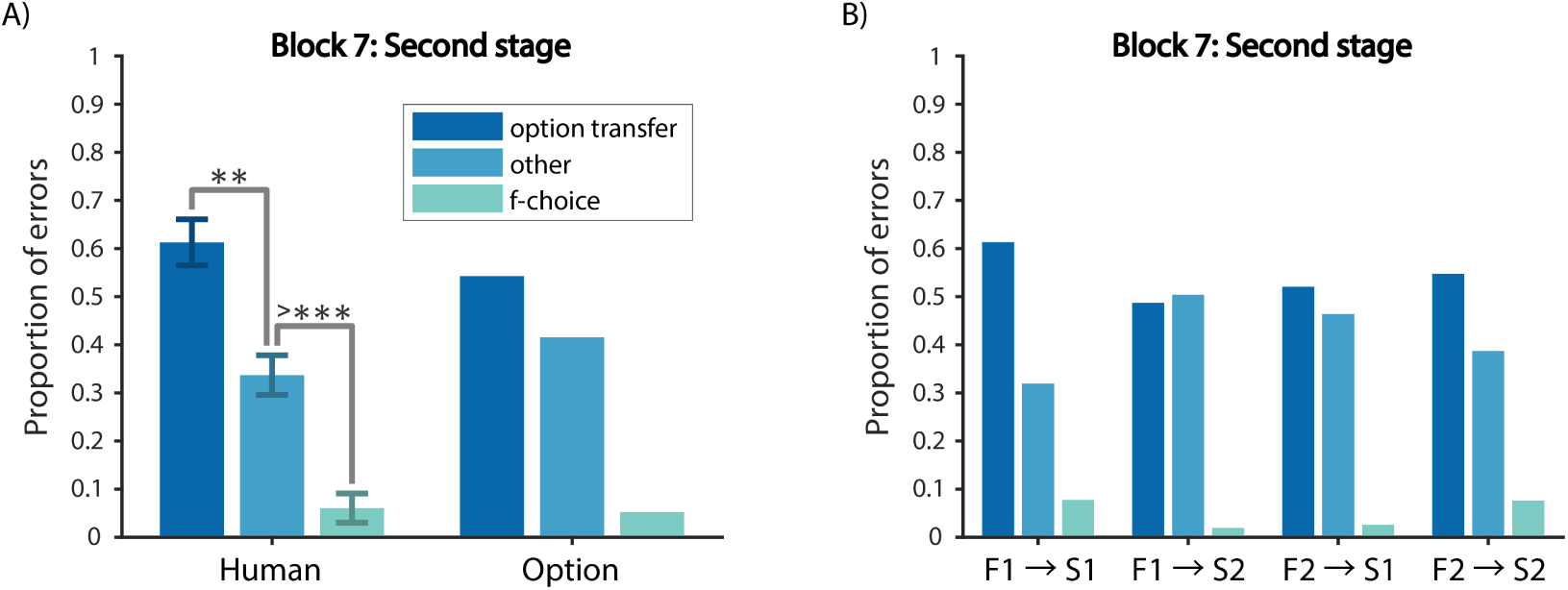
Experiment 2 second stage choices with parameters from Table 2 (A) Error type analysis of the second stage in Block 7 for participants (left) and the Option Model (right). (B) Error type analysis for each of the 4 branches in the second stage of Block 7 for the Option Model.

**Supplementary Figure S5:**
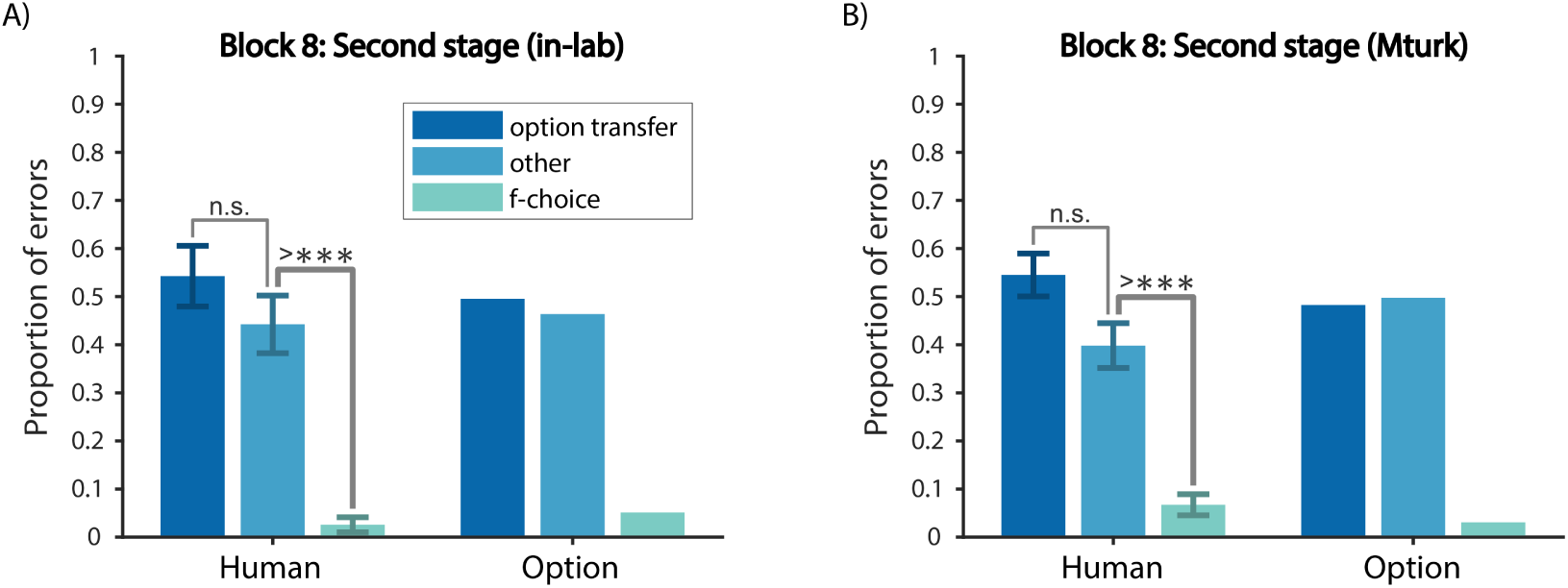
Experiment 3 second stage choices with parameters from Table 2. Error type analysis of the second stage in Block 8 for (A) in-lab participants (left) and the Option Model (right), and (B) Mturk participants (left) and the Option Model (right).

**Supplementary Figure S6:**
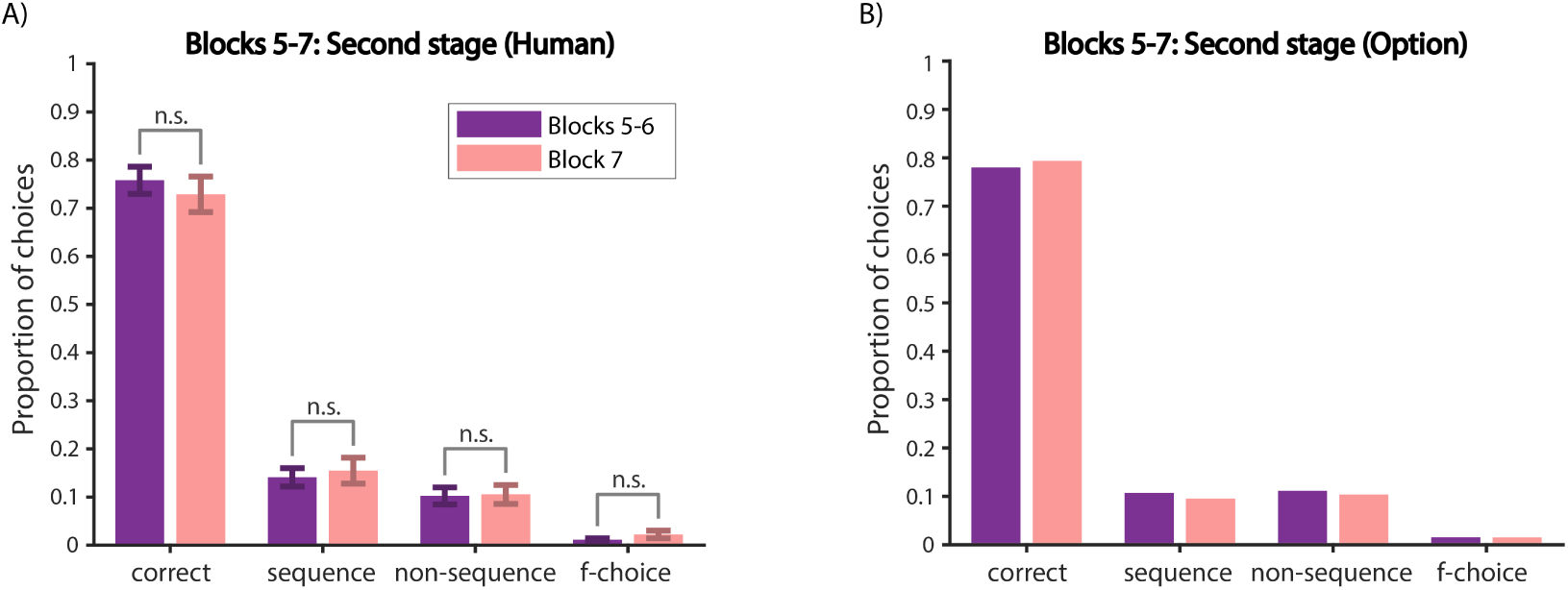
Experiment 1 second stage choices. Choice type analysis of the second stage comparing Blocks 5-6 and Block 7 for (A) participants and (B) the Option Mode. There was no significant difference across all choice types, indicating positive transfer in the second stage of Block 7.

**Supplementary Figure S7:**
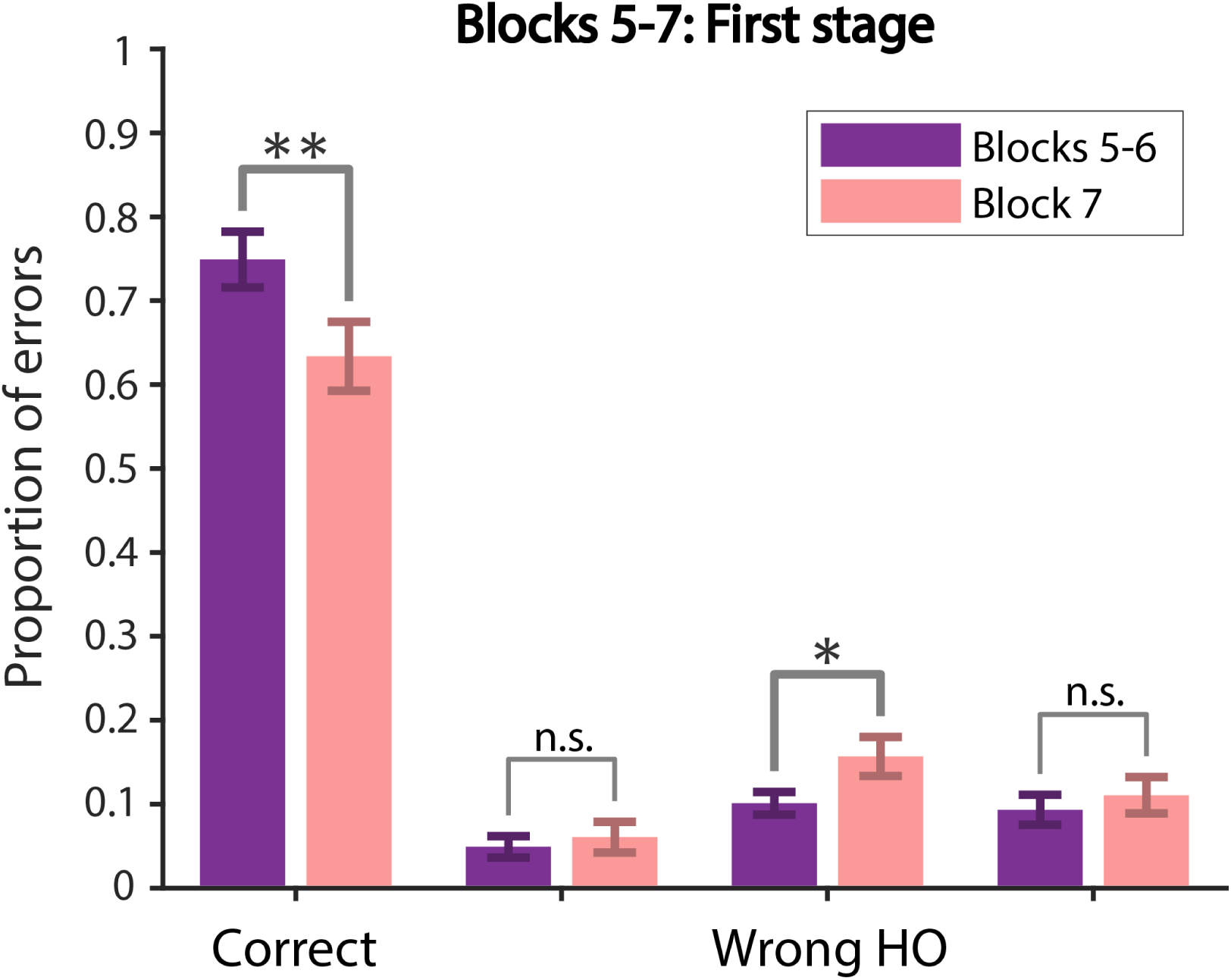
Experiment 2 first stage choices. Choice type analysis of the first stage comparing Blocks 5-6 and Block 7. The only error type that significantly increased was the wrong *HO* error, suggesting that participants were perseverating in the first stage while learning the new mappings in the second stage of Block 7.

**Supplementary Figure S8:**
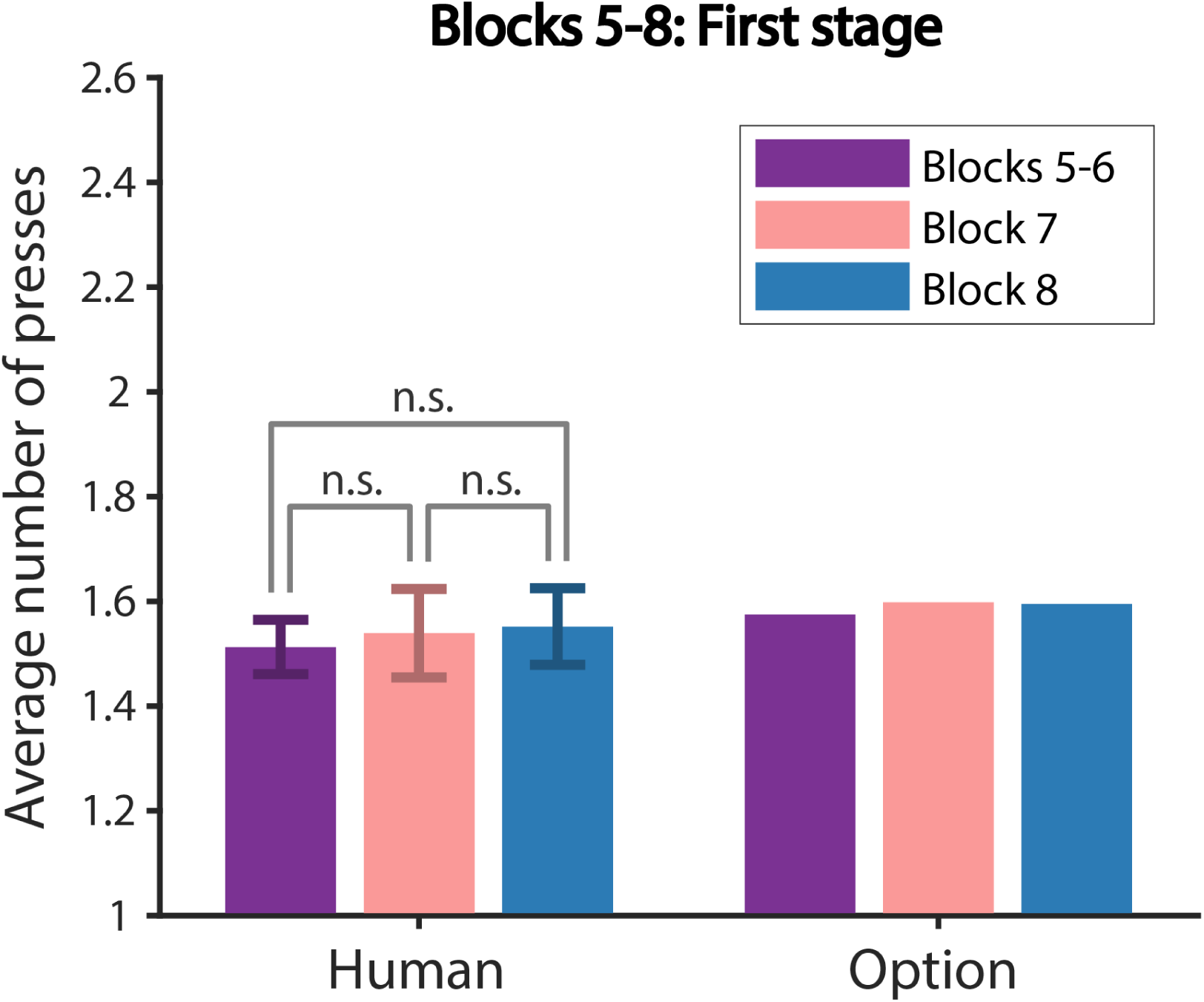
Experiment 3 Mturk first stage choices. Average number of presses in the first 10 trials of Blocks 5-8 in the first stage for participants (left) and the Option Model (right). This shows a lack of transfer in the first stage, representative of Experiments 3-4 first stage for both in-lab and Mturk populations.

**Supplementary Figure S9:**
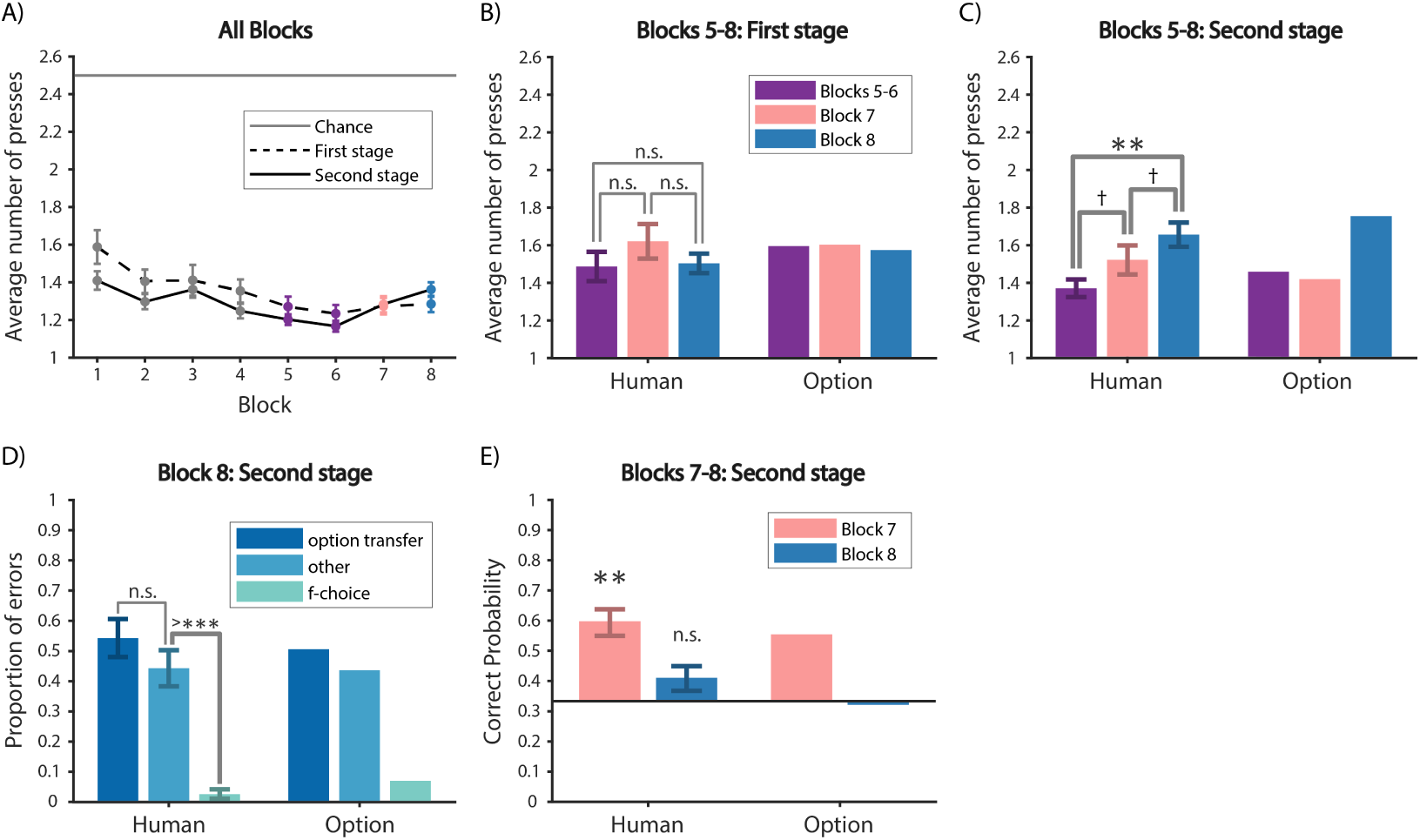
Experiment 3 summary. (A) Average number of key presses in the first and the second stages per block. (B) Average number of key presses for the first 10 trials of Blocks 5-8 for the first stage for participants (left) and the Option Model (right). (C) Same as (B) for the second stage. (D) Error type analysis of the second stage in Block 8 for participants (left) and the Option Model (right). The proportion of option transfer error was not significantly different from other error, different from Experiment 1 and Experiment 2, suggesting reduced option transfer. (E) Probability of a correct first key press for the second stage of the first trial of each of the 4 branches in Blocks 7-8 for participants (left) and the Option Model (right).

**Supplementary Figure S10:**
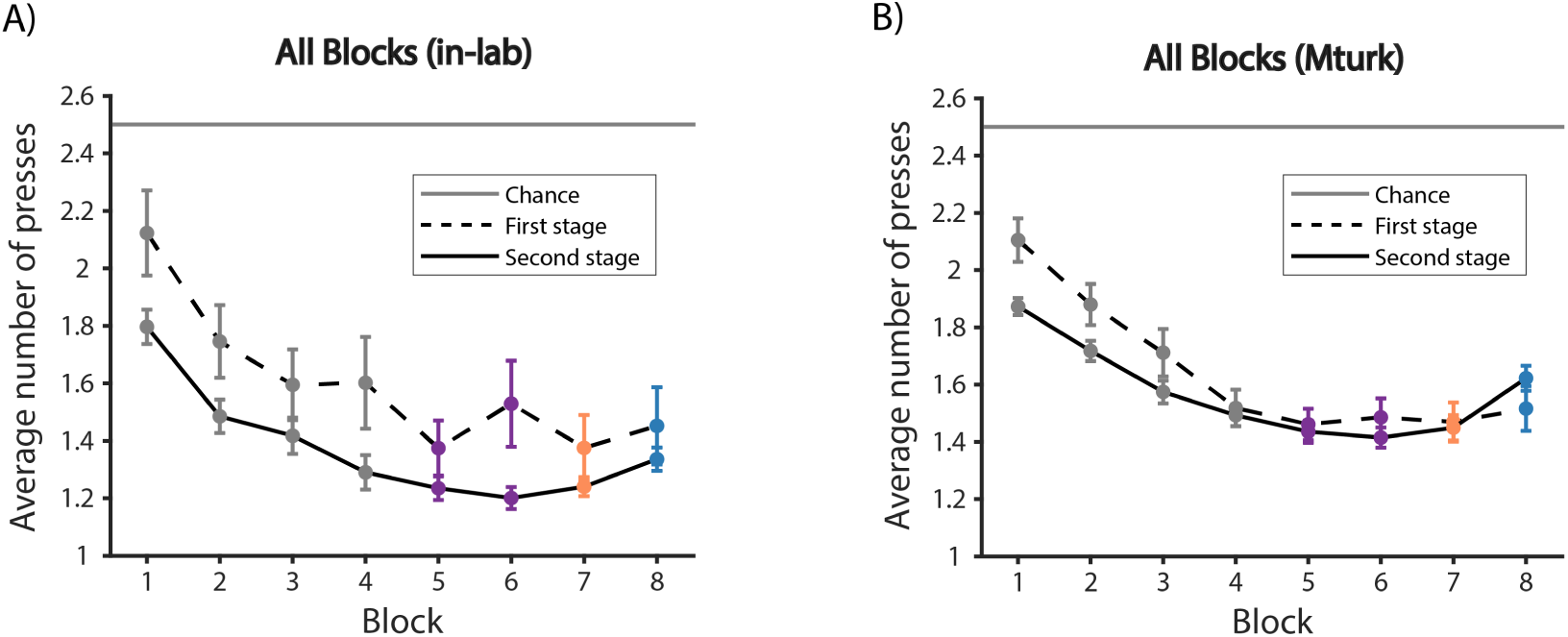
Experiment 4 number of presses. Average number of key presses in the first and the second stages per block for (A) in-lab participants and (B) Mturk participants.

**Supplementary Figure S11:**
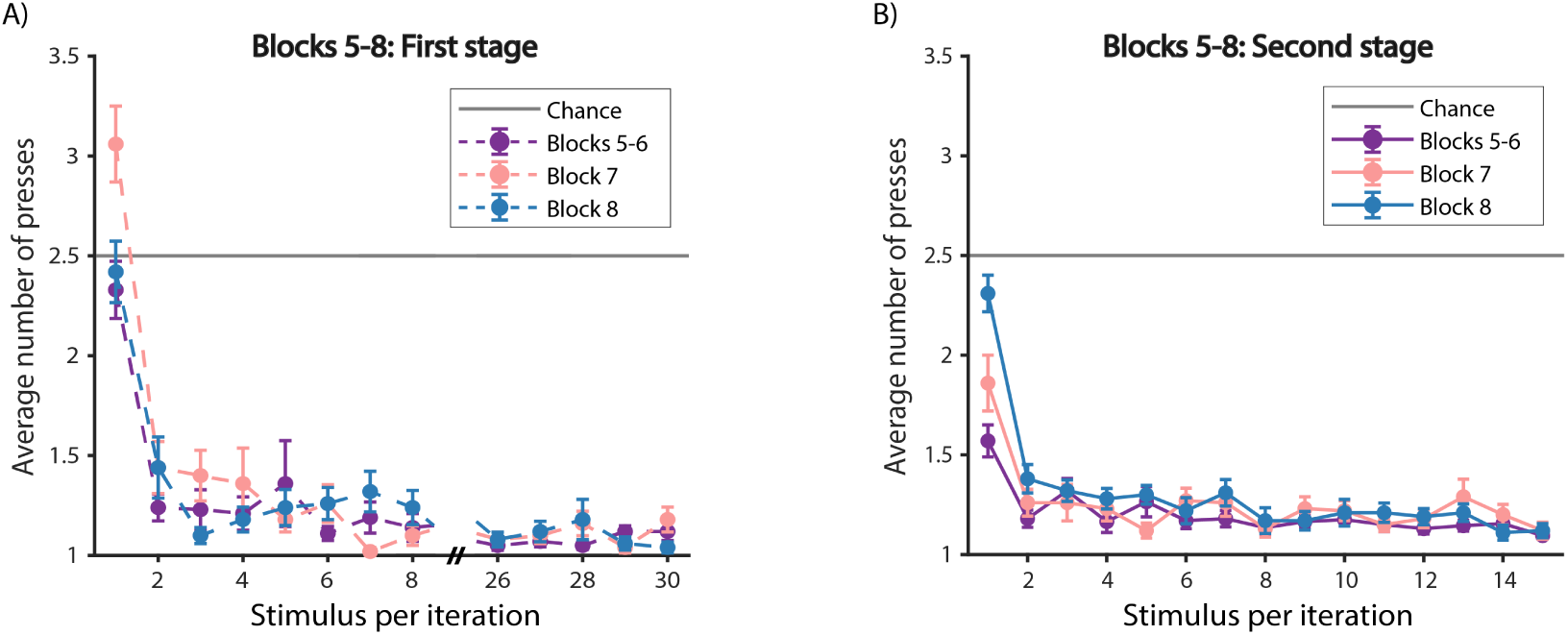
Experiment 1 performance within Blocks 5-8 for in-lab participants. (A) First stage. (B) Second stage.

**Supplementary Figure S12:**
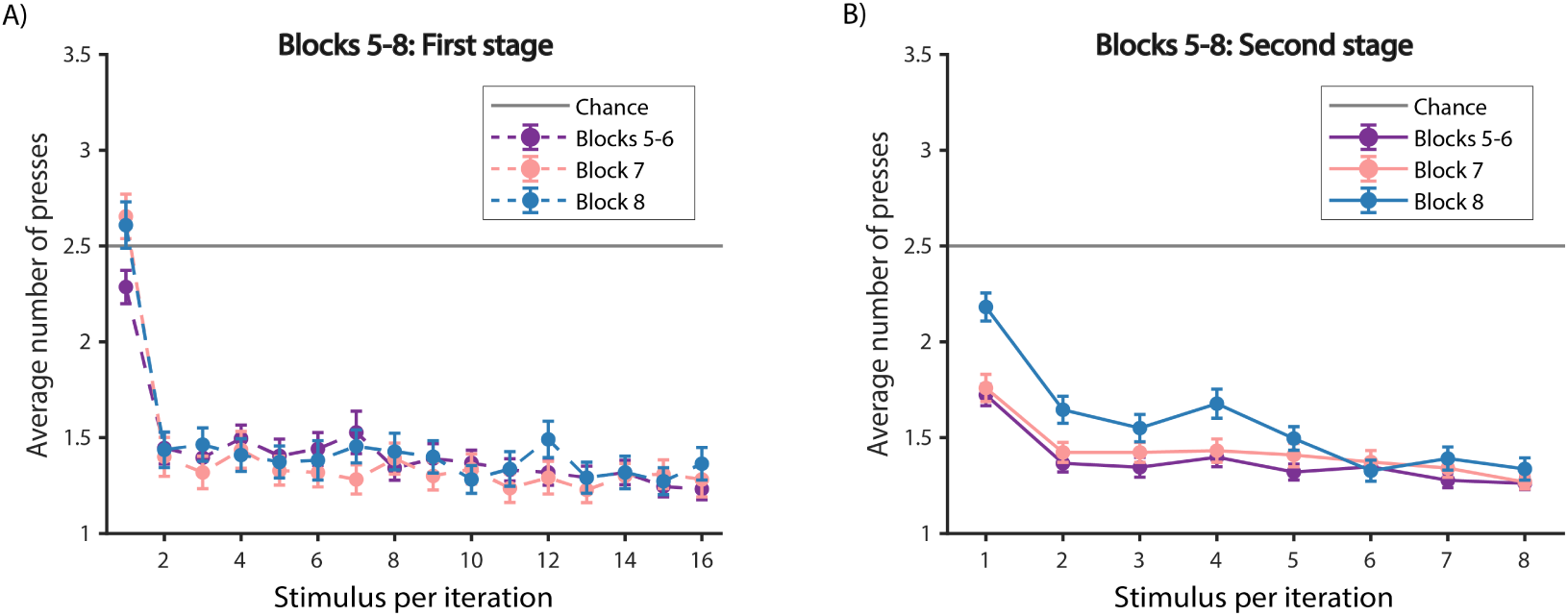
Experiment 1 performance within Blocks 5-8 for Mturk participants. (A) First stage. (B) Second stage.

**Supplementary Figure S13:**
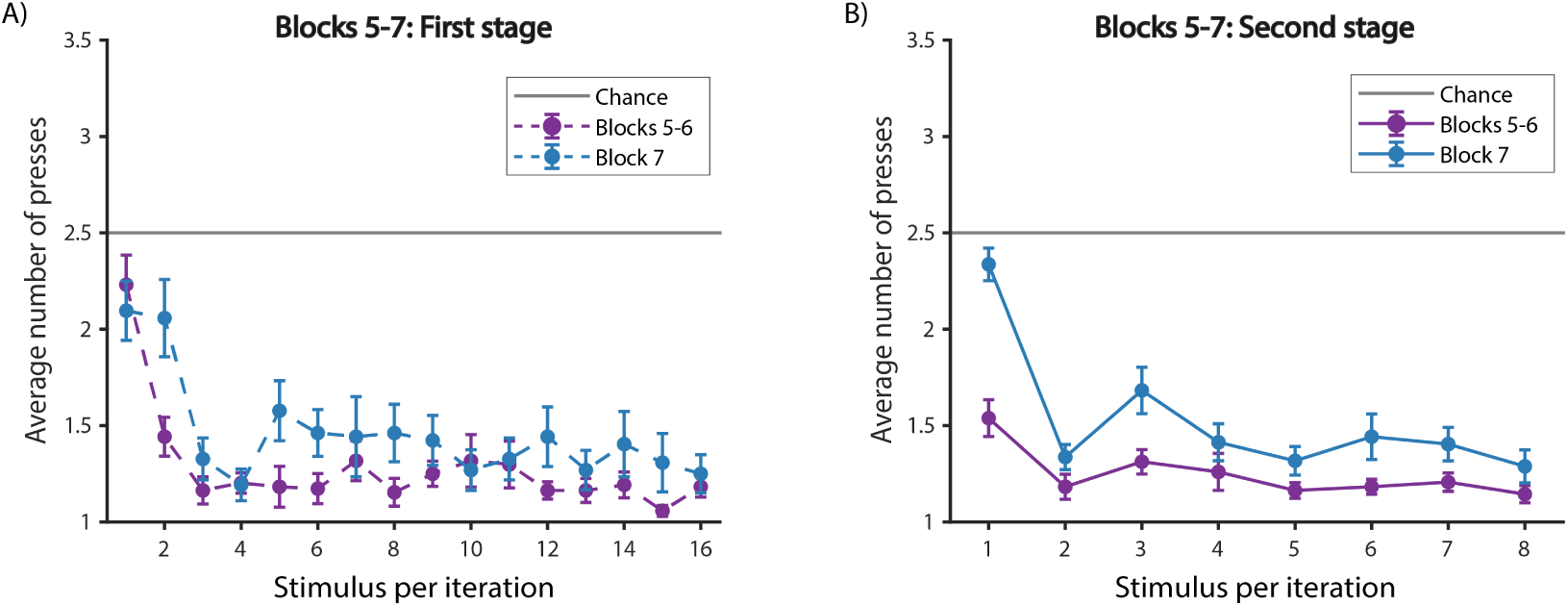
Experiment 2 performance within Blocks 5-7. (A) First stage. (B) Second stage.

**Supplementary Figure S14:**
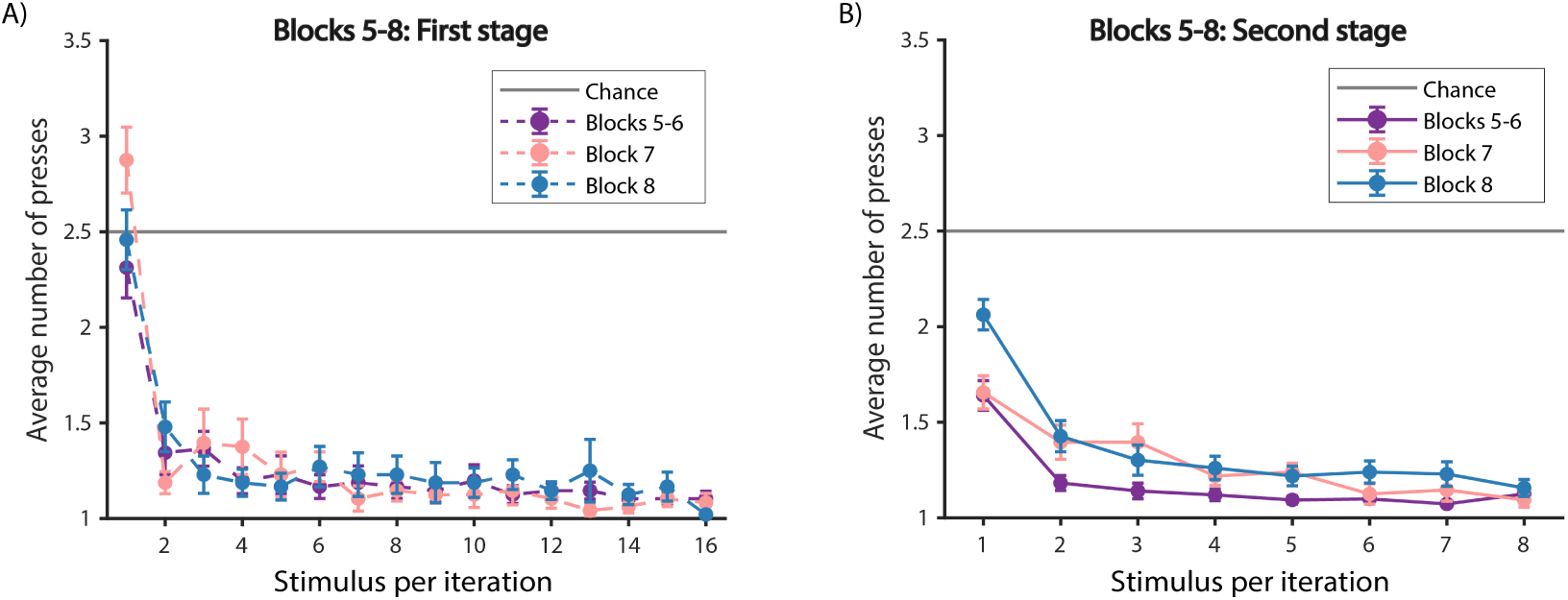
Experiment 3 performance within Blocks 5-8 for in-lab participants. (A) First stage. (B) Second stage.

**Supplementary Figure S15:**
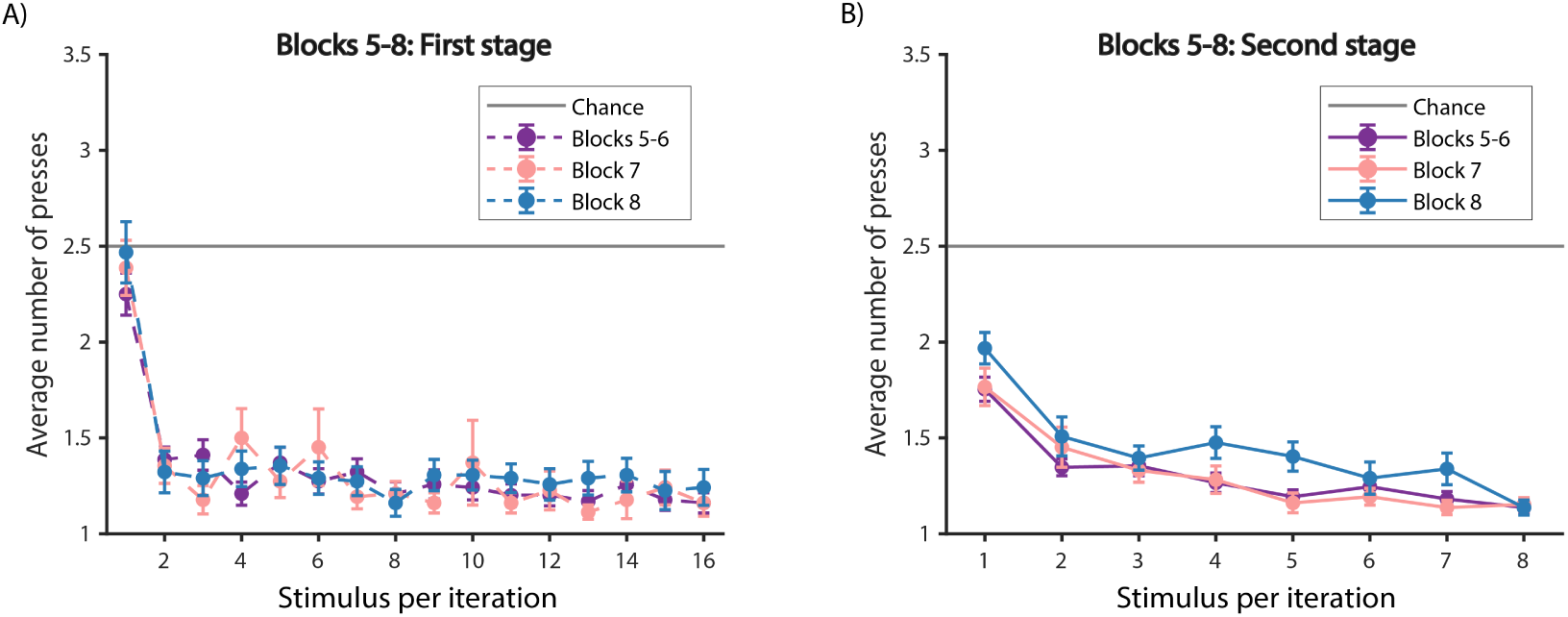
Experiment 3 performance within Blocks 5-8 for Mturk participants. (A) First stage. (B) Second stage.

**Supplementary Figure S16:**
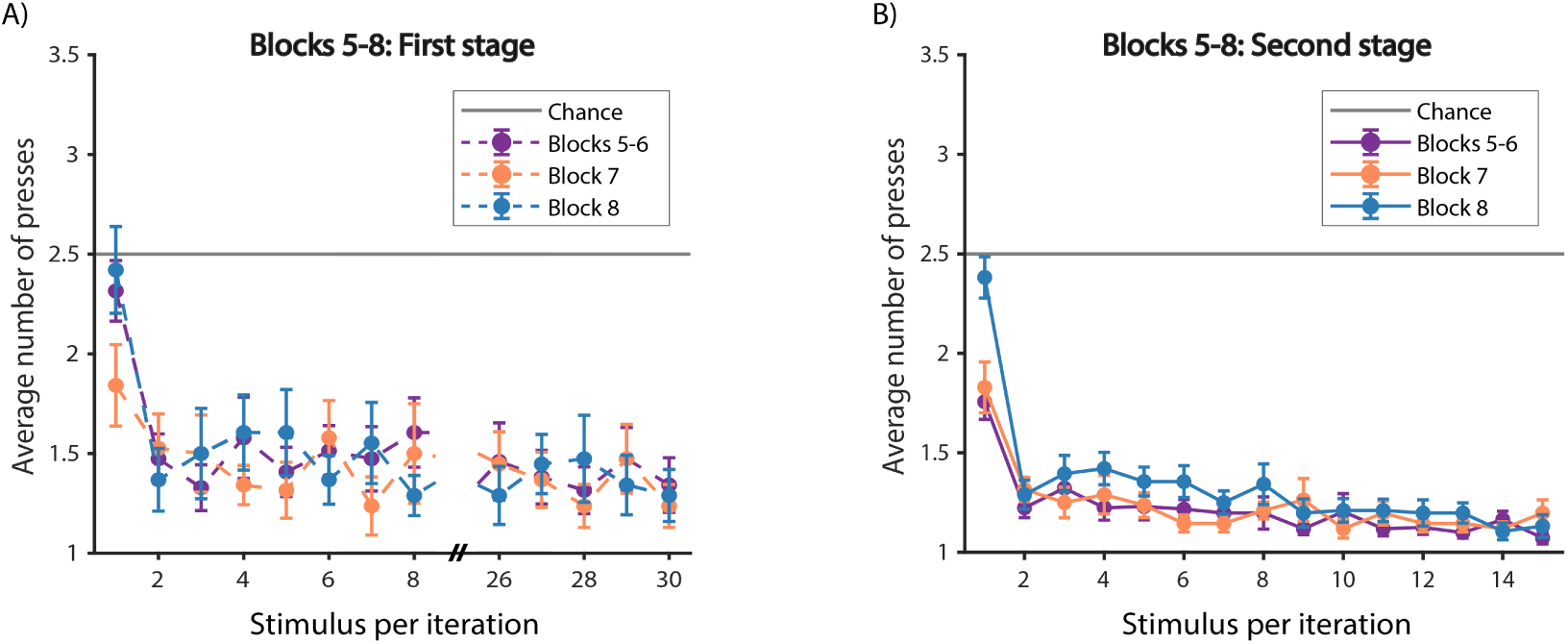
Experiment 4 performance within Blocks 5-8 for in-lab participants. (A) First stage. (B) Second stage.

**Supplementary Figure S17:**
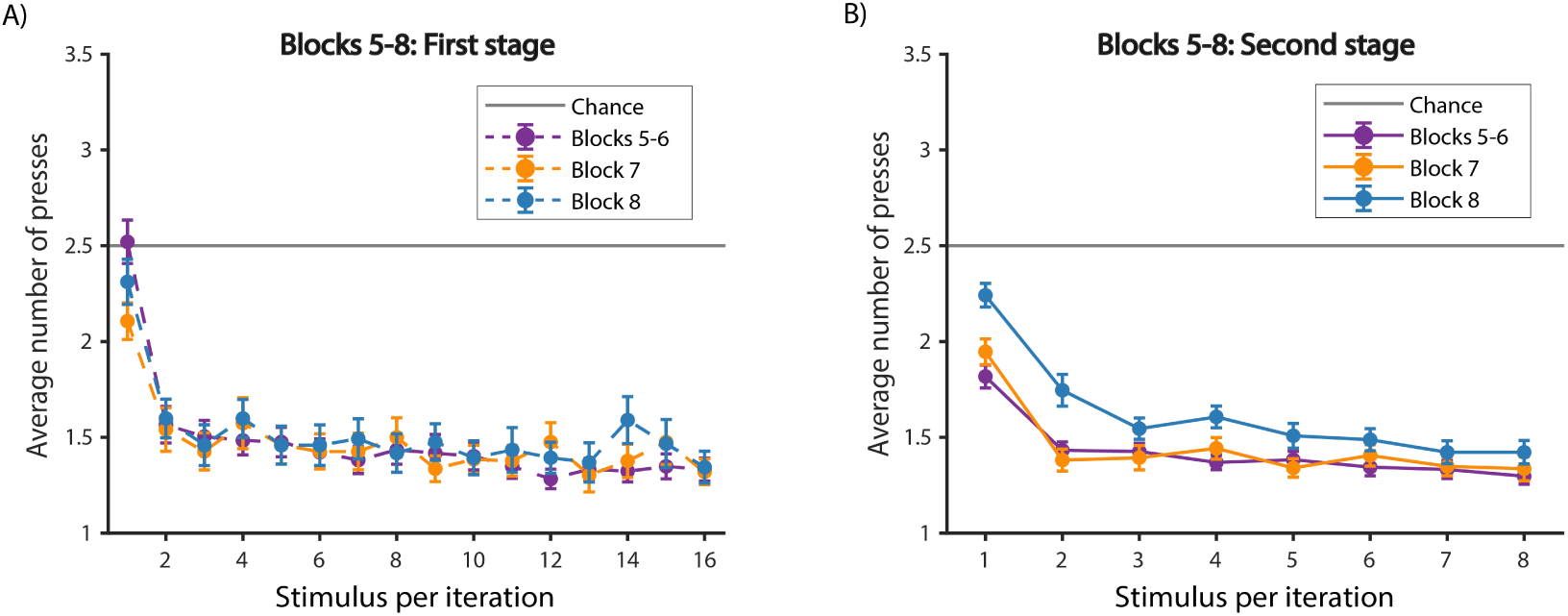
Experiment 4 performance within Blocks 5-8 for Mturk participants. (A) First stage. (B) Second stage.

## References

[1] R. S. Sutton, A. G. Barto, Reinforcement learning: An introduction, MIT press, 2018.

[2] V. Mnih, K. Kavukcuoglu, D. Silver, A. A. Rusu, J. Veness, M. G. Bellemare, A. Graves, M. Riedmiller, A. K. Fidjeland, G. Ostrovski, et al., Human-level control through deep reinforcement learning, Nature 518 (2015) 529.

[3] D. Silver, T. Hubert, J. Schrittwieser, I. Antonoglou, M. Lai, A. Guez, M. Lanctot, L. Sifre, D. Kumaran, T. Graepel, et al., A general reinforcement learning algorithm that masters chess, shogi, and go through self-play, Science 362 (2018) 1140–1144.

[4] Y. Niv, Reinforcement learning in the brain, Journal of Mathematical Psychology 53 (2009) 139–154.

[5] J. Gläscher, N. Daw, P. Dayan, J. P. O’Doherty, States versus rewards: dissociable neural prediction error signals underlying model-based and model-free reinforcement learning, Neuron 66 (2010) 585–595.

[6] Y. C. Leong, A. Radulescu, R. Daniel, V. DeWoskin, Y. Niv, Dynamic interaction between reinforcement learning and attention in multidimensional environments, Neuron 93 (2017) 451–463.

[7] S. Farashahi, K. Rowe, Z. Aslami, D. Lee, A. Soltani, Feature-based learning improves adaptability without compromising precision, Nature communications 8 (2017) 1768.

[8] A. G. Collins, M. J. Frank, How much of reinforcement learning is working memory, not reinforcement learning? a behavioral, computational, and neurogenetic analysis, European Journal of Neuroscience 35 (2012) 1024–1035.

[9] A. G. Collins, M. J. Frank, Cognitive control over learning: Creating, clustering, and generalizing task-set structure., Psychological review 120 (2013) 190.

[10] B. M. Lake, T. D. Ullman, J. B. Tenenbaum, S. J. Gershman, Building machines that learn and think like people, Behavioral and brain sciences 40 (2017).

[11] M. M. Botvinick, Y. Niv, A. C. Barto, Hierarchically organized behavior and its neural foundations: a reinforcement learning perspective, Cognition 113 (2009) 262–280.

[12] C. Diuk, A. Schapiro, N. Córdova, J. Ribas-Fernandes, Y. Niv, M. Botvinick, Divide and conquer: hierarchical reinforcement learning and task decomposition in humans, in: Computational and robotic models of the hierarchical organization of behavior, Springer, 2013, pp. 271–291.

[13] A. Solway, C. Diuk, N. Córdova, D. Yee, A. G. Barto, Y. Niv, M. M. Botvinick, Optimal behavioral hierarchy, PLoS computational biology 10 (2014) e1003779.

[14] E. Koechlin, C. Ody, F. Kouneiher, The architecture of cognitive control in the human prefrontal cortex, Science 302 (2003) 1181–1185.

[15] E. Koechlin, T. Jubault, Broca’s area and the hierarchical organization of human behavior, Neuron 50 (2006) 963–974.

[16] D. Badre, Cognitive control, hierarchy, and the rostro–caudal organization of the frontal lobes, Trends in cognitive sciences 12 (2008) 193–200.

[17] D. C. Van Essen, J. H. Maunsell, Hierarchical organization and functional streams in the visual cortex, Trends in neurosciences 6 (1983) 370–375.

[18] T. S. Lee, D. Mumford, Hierarchical bayesian inference in the visual cortex, JOSA A 20 (2003) 1434–1448.

[19] C. Wessinger, J. VanMeter, B. Tian, J. Van Lare, J. Pekar, J. P. Rauschecker, Hierarchical organization of the human auditory cortex revealed by functional magnetic resonance imaging, Journal of cognitive neuroscience 13 (2001) 1–7.

[20] J. Bill, H. Pailian, S. J. Gershman, J. Drugowitsch, Hierarchical structure is employed by humans during visual motion perception, bioRxiv (2019) 758573.

[21] N. Zarr, J. W. Brown, Hierarchical error representation in medial prefrontal cortex, NeuroImage 124 (2016) 238–247.

[22] O. Krigolson, C. Holroyd, Evidence for hierarchical error processing in the human brain, Neuroscience 137 (2006) 13–17.

[23] D. Badre, M. D’Esposito, Functional magnetic resonance imaging evidence for a hierarchical organization of the prefrontal cortex, Journal of cognitive neuroscience 19 (2007) 2082–2099.

[24] D. Badre, M. D’esposito, Is the rostro-caudal axis of the frontal lobe hierarchical?, Nature Reviews Neuroscience 10 (2009) 659.

[25] B. W. Balleine, A. Dezfouli, M. Ito, K. Doya, Hierarchical control of goal-directed action in the cortical–basal ganglia network, Current Opinion in Behavioral Sciences 5 (2015) 1–7.

[26] A. Dezfouli, B. W. Balleine, Actions, action sequences and habits: evidence that goal-directed and habitual action control are hierarchically organized, PLoS computational biology 9 (2013) e1003364.

[27] A. Dezfouli, B. W. Balleine, Habits, action sequences and reinforcement learning, European Journal of Neuroscience 35 (2012) 1036–1051.

[28] M. Tomov, S. Yagati, A. Kumar, W. Yang, S. Gershman, Discovery of hierarchical representations for efficient planning, BioRxiv (2018) 499418.

[29] M. K. Eckstein, A. G. Collins, Computational evidence for hierarchically-structured reinforcement learning in humans, bioRxiv (2019) 731752.

[30] M. J. Frank, D. Badre, Mechanisms of hierarchical reinforcement learning in corticostriatal circuits 1: computational analysis, Cerebral cortex 22 (2011) 509–526.

[31] D. Badre, M. J. Frank, Mechanisms of hierarchical reinforcement learning in cortico–striatal circuits 2: Evidence from fmri, Cerebral cortex 22 (2011) 527–536.

[32] A. G. Collins, J. F. Cavanagh, M. J. Frank, Human eeg uncovers latent generalizable rule structure during learning, Journal of Neuroscience 34 (2014) 4677–4685.

[33] M. Botvinick, D. C. Plaut, Doing without schema hierarchies: a recurrent connectionist approach to normal and impaired routine sequential action., Psychological review 111 (2004) 395.

[34] M. M. Botvinick, Multilevel structure in behaviour and in the brain: a model of fuster’s hierarchy, Philosophical Transactions of the Royal Society B: Biological Sciences 362 (2007) 1615–1626.

[35] A. G. Collins, Learning structures through reinforcement, in: Goal-Directed Decision Making, Elsevier, 2018, pp. 105–123.

[36] I. Biederman, Recognition-by-components: a theory of human image understanding., Psychological review 94 (1987) 115.

[37] B. M. Lake, R. Salakhutdinov, J. B. Tenenbaum, Human-level concept learning through probabilistic program induction, Science 350 (2015) 1332–1338.

[38] N. T. Franklin, M. J. Frank, Compositional clustering in task structure learning, PLoS computational biology 14 (2018) e1006116.

[39] D. Wingate, C. Diuk, T. O’Donnell, J. Tenenbaum, S. Gershman, Compositional policy priors (2013).

[40] J. Andreas, D. Klein, S. Levine, Modular multitask reinforcement learning with policy sketches, in: Proceedings of the 34th International Conference on Machine Learning-Volume 70, JMLR. org, pp. 166–175.

[41] D. Xu, S. Nair, Y. Zhu, J. Gao, A. Garg, L. Fei-Fei, S. Savarese, Neural task programming: Learning to generalize across hierarchical tasks, in: 2018 IEEE International Conference on Robotics and Automation (ICRA), IEEE, pp. 1–8.

[42] X. B. Peng, M. Chang, G. Zhang, P. Abbeel, S. Levine, Mcp: Learning composable hierarchical control with multiplicative compositional policies, arXiv preprint 1905.09808 (2019).

[43] R. S. Sutton, D. Precup, S. Singh, Between mdps and semi-mdps: A framework for temporal abstraction in reinforcement learning, Artificial intelligence 112 (1999) 181–211.

[44] M. Botvinick, A. Weinstein, Model-based hierarchical reinforcement learning and human action control, Phil. Trans. R. Soc. B 369 (2014) 20130480.

[45] A. McGovern, A. G. Barto, Automatic discovery of subgoals in reinforcement learning using diverse density (2001).

[46] I. Menache, S. Mannor, N. Shimkin, Q-cutdynamic discovery of subgoals in reinforcement learning, in: European Conference on Machine Learning, Springer, pp. 295–306.

[47] Ö. Şimşek, A. G. Barto, Using relative novelty to identify useful temporal abstractions in reinforcement learning, in: Proceedings of the twenty-first international conference on Machine learning, ACM, p. 95.

[48] M. C. Machado, C. Rosenbaum, X. Guo, M. Liu, G. Tesauro, M. Campbell, Eigenoption discovery through the deep successor representation, arXiv preprint 1710.11089 (2017).

[49] M. C. Machado, M. G. Bellemare, M. Bowling, A laplacian framework for option discovery in reinforcement learning, in: Proceedings of the 34th International Conference on Machine Learning-Volume 70, JMLR. org, pp. 2295–2304.

[50] Y. Jiang, S. Gu, K. Murphy, C. Finn, Language as an abstraction for hierarchical deep reinforcement learning, arXiv preprint 1906.07343 (2019).

[51] R. Fox, S. Krishnan, I. Stoica, K. Goldberg, Multi-level discovery of deep options, arXiv preprint 1703.08294 (2017).

[52] D. Jayaraman, F. Ebert, A. A. Efros, S. Levine, Time-agnostic prediction: Predicting predictable video frames, arXiv preprint 1808.07784 (2018).

[53] S. Nair, C. Finn, Hierarchical foresight: Self-supervised learning of long-horizon tasks via visual subgoal generation, arXiv preprint 1909.05829 (2019).

[54] D. Xu, R. Martín-Martín, D.-A. Huang, Y. Zhu, S. Savarese, L. Fei-Fei, Regression planning networks, arXiv preprint 1909.13072 (2019).

[55] J. F. Lehman, J. E. Laird, P. Rosenbloom, et al., A gentle introduction to soar, an architecture for human cognition, Invitation to cognitive science 4 (1996) 212–249.

[56] J. R. Anderson, D. Bothell, M. D. Byrne, S. Douglass, C. Lebiere, Y. Qin, An integrated theory of the mind., Psychological review 111 (2004) 1036.

[57] S. Nason, J. E. Laird, Soar-rl: Integrating reinforcement learning with soar, Cognitive Systems Research 6 (2005) 51–59.

[58] W.-T. Fu, J. R. Anderson, From recurrent choice to skill learning: A reinforcement-learning model., Journal of experimental psychology: General 135 (2006) 184.

[59] W. Schultz, P. Dayan, P. R. Montague, A neural substrate of prediction and reward, Science 275 (1997) 1593–1599.

[60] C. Diuk, K. Tsai, J. Wallis, M. Botvinick, Y. Niv, Hierarchical learning induces two simultaneous, but separable, prediction errors in human basal ganglia, Journal of Neuroscience 33 (2013) 5797–5805.

[61] J. J. Ribas-Fernandes, A. Solway, C. Diuk, J. T. McGuire, A. G. Barto, Y. Niv, M. M. Botvinick, A neural signature of hierarchical reinforcement learning, Neuron 71 (2011) 370–379.

[62] J. J. Ribas-Fernandes, D. Shahnazian, C. B. Holroyd, M. M. Botvinick, Subgoal-and goal-related reward prediction errors in medial prefrontal cortex, Journal of cognitive neuroscience 31 (2019) 8–23.

[63] A. C. Schapiro, T. T. Rogers, N. I. Cordova, N. B. Turk-Browne, M. M. Botvinick, Neural representations of events arise from temporal community structure, Nature neuroscience 16 (2013) 486.

[64] M. M. Botvinick, Hierarchical reinforcement learning and decision making, Current opinion in neurobiology 22 (2012) 956–962.

[65] C. B. Holroyd, N. Yeung, Motivation of extended behaviors by anterior cingulate cortex, Trends in cognitive sciences 16 (2012) 122–128.

[66] A. G. E. Collins, M. J. Frank, Neural signature of hierarchically structured expectations predicts clustering and transfer of rule sets in reinforcement learning, Cognition 152 (2016) 160–169.

[67] S. Monsell, Task switching, Trends in cognitive sciences 7 (2003) 134–140.

[68] J. Pitman, Combinatorial Stochastic Processes: Ecole d’Eté de Probabilités de Saint-Flour XXXII-2002, Springer, 2006.

[69] J. X. Wang, Z. Kurth-Nelson, D. Kumaran, D. Tirumala, H. Soyer, J. Z. Leibo, D. Hassabis, M. Botvinick, Prefrontal cortex as a meta-reinforcement learning system, Nature neuroscience 21 (2018) 860.

[70] A. G. E. Collins, M. J. Frank, Motor demands constrain cognitive rule structures, PLoS computational biology 12 (2016) e1004785.

[71] M. Sarafyazd, M. Jazayeri, Hierarchical reasoning by neural circuits in the frontal cortex, Science 364 (2019) eaav8911.

[72] R. Moran, M. Keramati, P. Dayan, R. J. Dolan, Retrospective model-based inference guides model-free credit assignment, Nature communications 10 (2019) 750.

[73] B. A. Clegg, G. J. DiGirolamo, S. W. Keele, Sequence learning, Trends in cognitive sciences 2 (1998) 275–281.

[74] G. Konidaris, A. G. Barto, Building portable options: Skill transfer in reinforcement learning., in: IJCAI, volume 7, pp. 895–900.

[75] K. R. Allen, K. A. Smith, J. B. Tenenbaum, The tools challenge: Rapid trial-and-error learning in physical problem solving, arXiv preprint 1907.09620 (2019).

[76] A. G. Collins, The cost of structure learning, Journal of Cognitive Neuroscience 29 (2017) 1646–1655.

[77] J. Y. Angela, J. D. Cohen, Sequential effects: superstition or rational behavior?, in: Advances in neural information processing systems, pp. 1873–1880.

[78] K. L. Stachenfeld, M. M. Botvinick, S. J. Gershman, The hippocampus as a predictive map, Nature neuroscience 20 (2017) 1643.

[79] I. Momennejad, E. M. Russek, J. H. Cheong, M. M. Botvinick, N. D. Daw, S. J. Gershman, The successor representation in human reinforcement learning, Nature Human Behaviour 1 (2017) 680.

